# Mechanism of small molecule inhibition of *Plasmodium falciparum* myosin A informs antimalarial drug design

**DOI:** 10.1101/2022.09.09.507123

**Authors:** Dihia Moussaoui, James P. Robblee, Daniel Auguin, Fabio Fisher, Patricia M. Fagnant, Jill E. Macfarlane, Julia Schaletzky, Eddie Wehri, Christoph Mueller-Dieckmann, Jake Baum, Julien Robert-Paganin, Kathleen M. Trybus, Anne Houdusse

**Affiliations:** Structural Motility, Institut Curie, Université Paris Sciences et Lettres, Sorbonne Université, CNRS UMR144, 75248 Paris, France; Department of Molecular Physiology & Biophysics, University of Vermont, Burlington, VT, USA; Laboratoire de Biologie des Ligneux et des Grandes Cultures (LBLGC), Université d’Orléans, INRAE, USC1328, Orléans, France; Department of Life Sciences, Imperial College London, Exhibition Road, South Kensington, London SW7 2AZ, UK; School of Medical Sciences, Faculty of Medicine & Health, UNSW Sydney, Kensington, 2052, NSW, Australia; Structural biology group, European Synchrotron Radiation Facility. 71, Avenue des Martyrs, 38000 Grenoble, France. Current affiliation of DM; Center for Emerging and Neglected Diseases, Drug Discovery Center, Berkeley, CA, USA

**Author notes:** Correspondence and requests for materials should be addressed to K.M.T. or to A.H. co-first authors (Dihia Moussaoui, James P. Robblee). **Data availability** The atomic models of PfMyoA/Apo and PfMyoA/KNX-002 are available on the PDB under the accession codes 8A12 and 8A13 respectively. **Author Contributions** AH and KMT designed and directed the research. DM crystallized PfMyoA and PfMyoA complexed to KNX-002. DM and CMD collected the data. DM processed the data, solved and refined the structures. DA performed the molecular dynamics calculations. JPR performed the *in vitro* functional assays. PMF and JEM expressed and purified PfMyoA. FF performed the parasitemia assays. JS and EW developed the screening assay and performed the high-throughput screen and hit characterization for CK2140597/KNX-002 while at Cytokinetics. DM, JPR, DA, JRP, KMT and AH analyzed and discussed the results. DM, JRP and AH wrote the initial version of the paper, with the help of JPR, DA, JB and KMT. All the authors were involved in reviewing and editing the paper. JB, JRP, KMT and AH were involved in project administration.

**Keywords:** Malaria, *Plasmodium falciparum*, Myosin A, KNX-002, antimalarial treatment

## Abstract

Malaria is responsible for more than a half million deaths per year. The *Plasmodium* parasites responsible continue to develop resistance to all known agents, despite treatment with different antimalarial combinations. The atypical Myosin A motor (PfMyoA) is part of a core macromolecular complex called the glideosome, essential for *Plasmodium* parasite mobility and therefore an attractive drug target. Here, we characterize the interaction of a small molecule (KNX-002) with PfMyoA. KNX-002 inhibits PfMyoA ATPase activity *in vitro* and blocks asexual blood stage growth of merozoites, one of three motile *Plasmodium* life-cycle stages. Combining biochemical assays, X-ray crystallography and molecular dynamics, we demonstrate that KNX-002 targets a novel pocket in PfMyoA, sequestering it in a post-rigor state detached from actin. KNX-002 binding affects Mg^2+^ coordination near ATP, preventing ATP hydrolysis and thus inhibiting motor activity. This first-in-class small-molecule inhibitor of PfMyoA paves the way for developing a new generation of antimalarial treatments.

## Introduction

Malaria infection in humans, caused by unicellular parasites from the genus *Plasmodium* and transmitted via the bite of an infected *Anopheles* mosquito, is a major global health challenge^1,2^. 627,000 people died of malaria in 2020, the majority being children under the age of 5 years^3^. Despite the remarkable progress of research aimed at advancing antimalarial therapeutics, the parasite continues to become resistant to all existing treatments, including the gold-standard front line artemisinin-based combination therapies (ACTs) ^4,5^. Recent licensure of the first ever malaria vaccine heralds a new era in efforts to control malaria, but the relatively modest efficacy of the RTS,S vaccine means that complementary approaches will be essential if the WHO’s goal of a 90% reduction in rates by 2030 is to be realized^6^.

Malaria parasites are motile throughout their complex human and mosquito life cycle. They move by a process called gliding motility^7^, which underpins their ability to reach, cross, and enter host tissues and cells. Gliding is powered by a macromolecular complex called the glideosome, the core of which is comprised of a divergent class XIV myosin A (PfMyoA) interacting with short, oriented filaments of a divergent actin (PfAct1)^7,8^. Importantly, PfMyoA has been demonstrated to be essential for parasite motility and for pathogenesis^9,10,11,12^, thus making this myosin motor a desirable target for preventing lifecycle progression and as such malaria disease.

The development of small molecules able to specifically activate or inhibit myosin force production has been successful in several other myosin classes, including the first-in-class activator Omecamtiv mecarbil (OM) and the first-in-class inhibitor Mavacamten, targeting β-cardiac myosin against heart failure and inherited cardiac diseases^13,14,15^; MPH-220, an inhibitor of skeletal muscle myosin (SkMyo2) against muscular spasticity^16^; and CK-571, a smooth muscle myosin 2 inhibitor (SmMyo2) against asthma^17^; reviewed by ^18,19,20^. The fact that some of these compounds such as OM and Mavacamten have completed phase 3 clinical trials ^21,22^ gives significant confidence to the development of myosins as realistic and druggable targets. Mavacamten has recently been approved by the FDA (trade name Camzyos) for the treatment of obstructive hypertrophic cardiomyopathy to improve functional capacity and symptoms.

Here, we characterize the interaction of PfMyoA with KNX-002, a small molecule inhibitor of PfMyoA ATPase activity and of *P. falciparum* merozoite invasion of red blood cells. The X-ray structure of PfMyoA complexed to KNX-002 identifies the binding pocket, and combined with molecular dynamics, transient kinetics and binding studies reveal how the compound sequesters PfMyoA in a state of low affinity for actin. Our results demonstrate that KNX-002 is a promising candidate for the development of novel antimalarial treatment.

## Results

### KNX-002 is a promising first-in-class small molecule antimalarial inhibitor

KNX-002 was initially identified as an inhibitor of PfMyoA from high throughput actin-activated ATPase screens (see Methods). It inhibits both the actin activated (IC_50_ = 7.2 μM) and the basal (IC_50_ = 3.6 μM) ATPase activity of PfMyoA, while showing little effect on cardiac or skeletal myosin II (**Fig. 1a**). Measurement of the affinity of KNX-002 for nucleotide-free PfMyoA (8.6 μM) and PfMyoA.ADP (11.1 μM) yielded values (**Supplementary Fig. 1**) that were of the same order of magnitude as the IC_50_ obtained from the steady-state ATPase activities. KNX-002 also inhibited the ability of PfMyoA to move actin in an *in vitro* motility assay. In the absence of KNX-002 robust motility of actin filaments was observed at speeds of 3.33 ± 0.50 μm/s (**Supplementary Movie 1**), while there was no directed actin filament motion in the presence of 200 μM KNX-002 (**Supplementary Movie 2**). Very few filaments were seen in the presence of KNX-002 (6.5 ± 1.9 versus 110.3 ± 1.9 filaments/field, n=6 fields).

**Figure 1.**
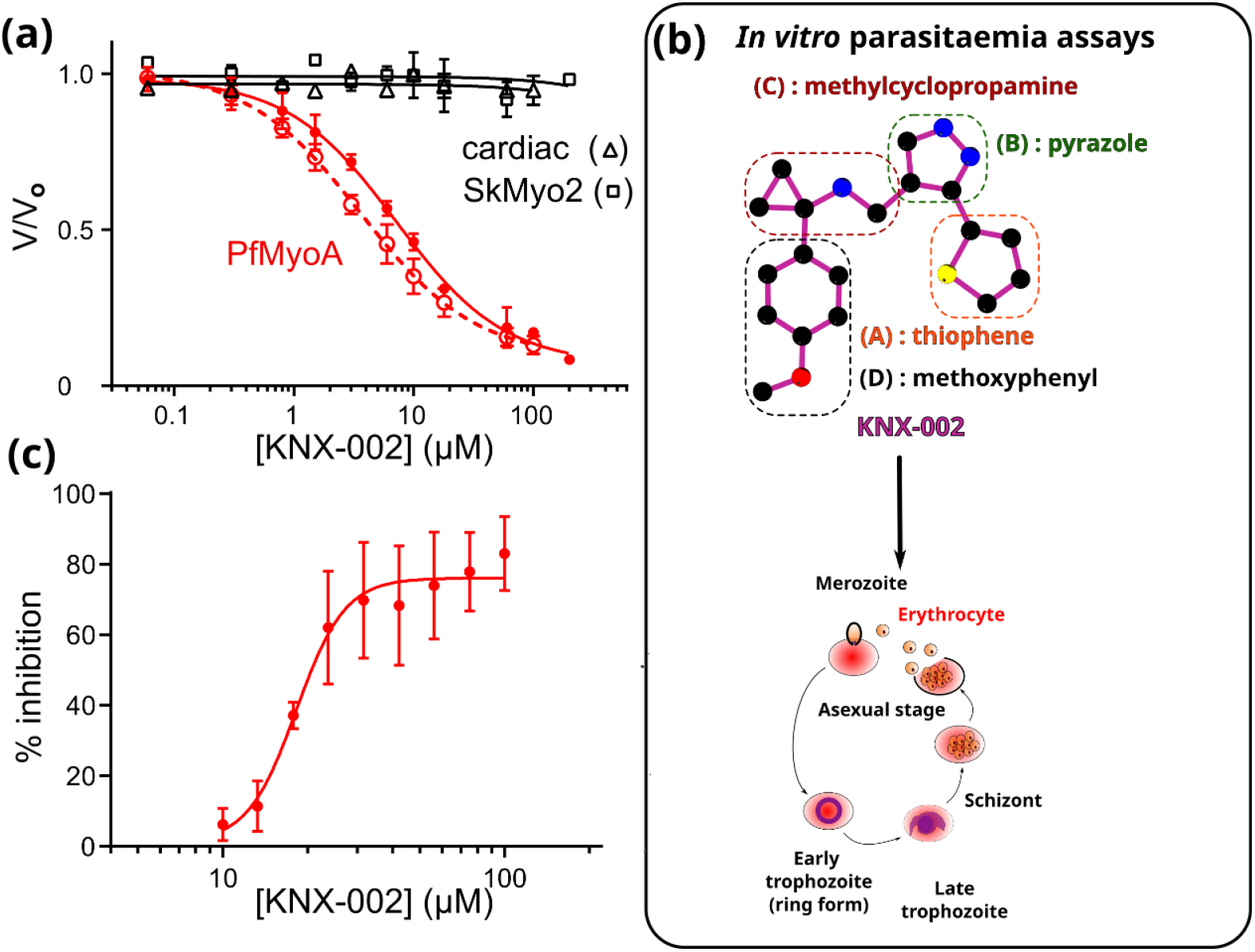
KNX-002 inhibits PfMyoA. **(a)** KNX-002 inhibits actin-activated (filled red circles, IC_50_=7.2 μM, (95% CI, 5.8-9.0 μM), n=3) and basal ATPase activity (open red circles, IC_50_=3.6 μM (95% CI, 3.0-4.5 μM), n=3). KNX-002 has little effect on the actin-activated ATPase of skeletal myosin (SkMyo2, black squares, IC_50_ > 200 μM, n=2) or cardiac myosin (black triangles, IC_50_ > 100 μM, n=2). **(b)** The constituent groups of the KNX-002 structure are indicated. **(c)** The inhibition of red blood cell invasion by KNX-002 on asexual, blood-stage growth was quantified (IC_50_ = 18.2 μM (95% CI, 15.7-22.3 μM)).

Because PfMyoA is essential for blood cell invasion ^9,10^, we tested the effect of KNX-002 on *P. falciparum* asexual, blood-stage growth, itself dependent on the ability of merozoites to invade erythrocytes ^23^. KNX-002 inhibited asexual blood stage growth of merozoites (IC_50_= 18.2 μM), confirming a drug effect on parasite cells *ex vivo* (**Fig. 1b, 1c**). Taken together, these results demonstrate that KNX-002 is an inhibitor of PfMyoA activity *in vitro*, as well as impeding asexual parasite blood stage growth that is a measure of the ability of parasites to invade, replicate, exit and reinvade erythrocytes, resulting in the symptomatic stage of malaria.

The effect of KNX-002 on the growth of three human cell lines (MRC5 fibroblasts, RPE1 epithelial and HepG2 hepatic) was evaluated (**Supplementary Table 1**). 20 μM KNX-002 showed <21% growth inhibition of HepG2 cells, and no measurable inhibition of the other two cell types, suggesting that compounds with similar properties could be promising hits to develop deliverable treatments.

### KNX-002 targets the post-rigor state

To dissect the inhibition mechanism of KNX-002 on the force production cycle, we crystallized full-length PfMyoA complexed with KNX-002. We successfully solved the structure of full-length PfMyoA complexed to both the ATP analogue Mg.ATP-gamma-S (Mg.ATPγS) and KNX-002 (PfMyoA/KNX-002) at a resolution of 2.14 Å. The structure of full-length PfMyoA complexed to Mg.ATPγS without the compound (PfMyoA/Apo) was also solved at a resolution of 2.03 Å (**Supplementary Table 2**).

These experiments allow a direct comparison of the apo and the KNX-002-bound structures when a hydrolysable ATP analog is bound. The high-resolution electron density maps are at similar resolution in these two datasets and allow us to build the nucleotides and KNX-002 without ambiguity, as well as the water molecules, in particular those in the active site (**Fig. 2a, 2b**). Whether bound to KNX-002 or not, PfMyoA crystallized in the post-rigor (PR) state, an ATP-bound myosin structural state with low affinity for the actin track which is populated upon detachment of the motor from the track prior to the priming of its lever arm (**Supplementary Fig. 2**). The two structures are highly superimposable (rmsd 0.214 Å on 914 Cα-atoms, **Fig. 2c**), indicating that KNX-002 does not induce major structural changes in the myosin structure. Some limited adjustments occur for the residues around the pocket upon KNX-002 binding (max difference in Cα positions being <2 Å, **Supplementary Fig. 3a**). The compound is close to the active site and Switch-2, an important connector of the motor that changes its conformation during the recovery stroke to favor ATP hydrolysis (**Supplementary Fig. 2**).

**Figure 2.**
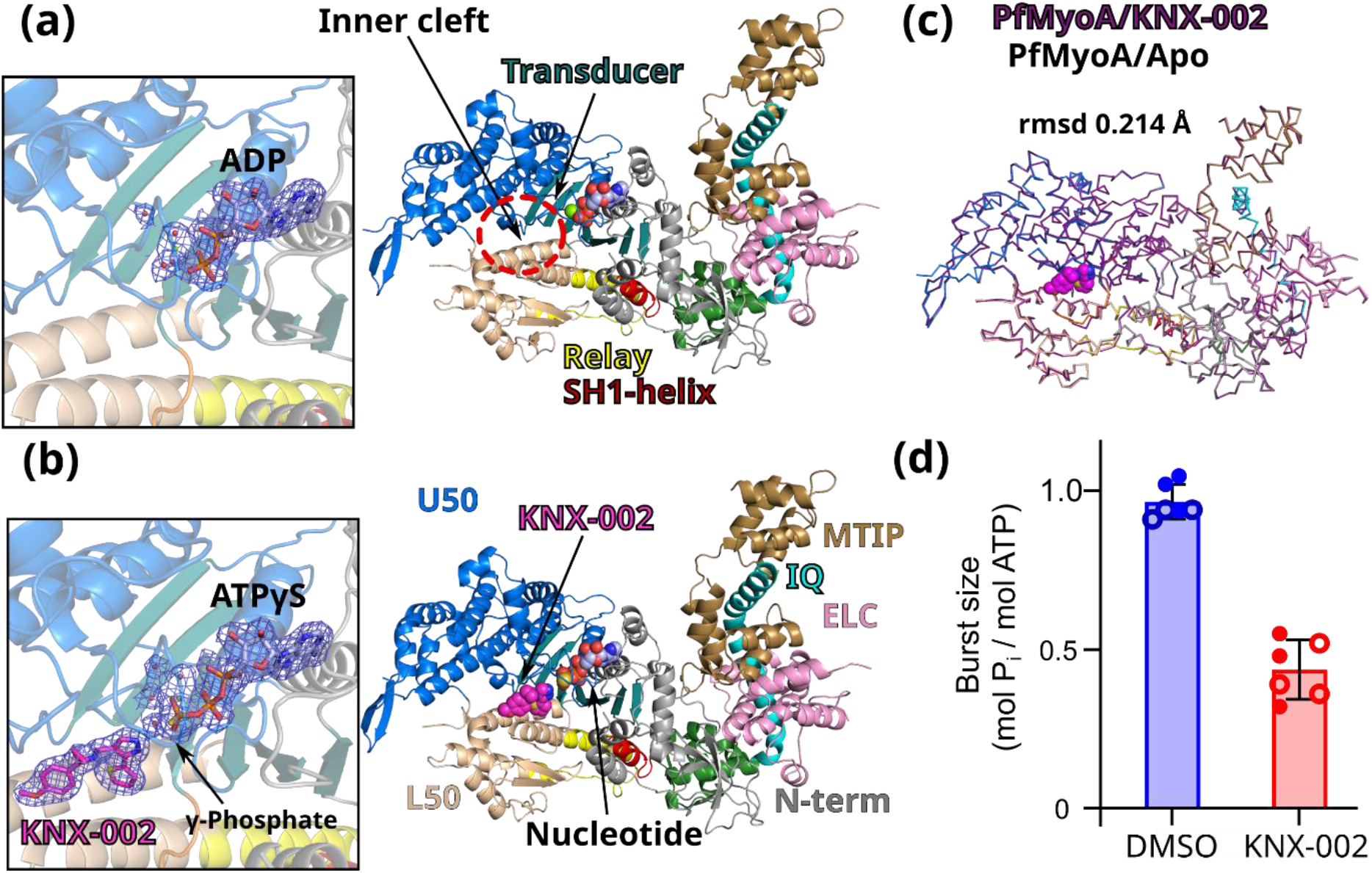
KNX-002 targets the post-rigor (PR) state. **(a)** Structure of PfMyoA in the apo condition (PfMyoA/Apo). Only ADP is found in the 2Fo-Fc electron density map contoured at 1.0 σ (on the left). ATPyS was thus hydrolyzed by myosin, in contrast to the same experiment performed in the presence of KNX-002 **(b)** Structure of PfMyoA complexed with KNX-002 and MgATPyS (PfMyoA/KNX-002). The compound, MgATPyS, the water molecules and the Mg^2+^ ion can be clearly identified in the 2Fo-Fc electron density map contoured at 1.0 σ (on the left). The gamma-phosphate of ATPyS is present in the density. **(c)** PfMyoA/KNX-002 and PfMyoA/Apo superimpose with a rmsd of 0.215 Å on the Cα and are both in a PR state. The compound does not induce major structural rearrangements upon binding. **(d)** Manual quenching experiments show a decreased phosphate burst from 0.97 ± 0.05 mol P_i_/ mol ATP in the absence of compound to 0.44 ± 0.08 mol P_i_/ mol ATP in the presence of 100 μM KNX-002 (p<0.0001, two-tailed t-test with Welch’s correction). Data represent two experiments each performed in triplicate with independent protein preparations (open and filled circles).

The major difference between the apo and KNX-002-bound structures is the state and coordination of the nucleotide. While unhydrolyzed Mg.ATPyS occupies the active site of PfMyoA/KNX-002, the substrate was hydrolyzed in PfMyoA/Apo and only Mg.ADP remains in the active site (**Fig. 2a, 2b**). Because cleavage was only observed in the absence of compound, this result implies that KNX-002 directly inhibits the hydrolysis process.

To test this hypothesis, we performed manual quench experiments. In the presence of KNX-002, the phosphate burst is reduced 2.2-fold (**Fig. 2d**), which demonstrates the ability of KNX-002 to directly reduce ATP hydrolysis. In the motor cycle, ATP hydrolysis occurs after the recovery stroke. This ATP-bound transition is essential for lever arm priming and leads to the pre-powerstroke state (PPS) in which nucleotide-binding loops, notably Switch-2, are positioned to favor ATP hydrolysis (**Supplementary Fig. 2**). Our results suggest that KNX-002 inhibits hydrolysis by stabilizing the Post-rigor state, thus blocking the recovery stroke. This agrees with our observation that attempts at crystallizing the motor in the presence of KNX-002 when the motor adopts the pre-powerstroke state (see Methods) lead to crystals lacking compound. In fact, the conformation of the KNX-002 binding pocket is drastically different in the PPS state and thus becomes incompetent for KNX-002 binding (**Supplementary Fig. 3b**).

### KNX-002 binds in a previously undescribed pocket

In the structure, KNX-002 is buried in a tight pocket located between the U50 and L50 subdomains, in the “inner cleft” (**Fig. 2a, 3a**), close to the so-called back-door of myosin ^24,25^, located near the yP_i_ of the nucleotide (**Fig. 3a**). Specific connectors essential in allosteric transduction ^25,26^ are directly involved in the binding of KNX-002: Switch-1, Switch-2 and Transducer (**Fig. 3a**). This structure reveals the first example of a small-molecule co-crystallized in the PR state while binding near Switch-2. Most compounds described to date target the PPS state ^14,27,16^.

**Figure 3.**
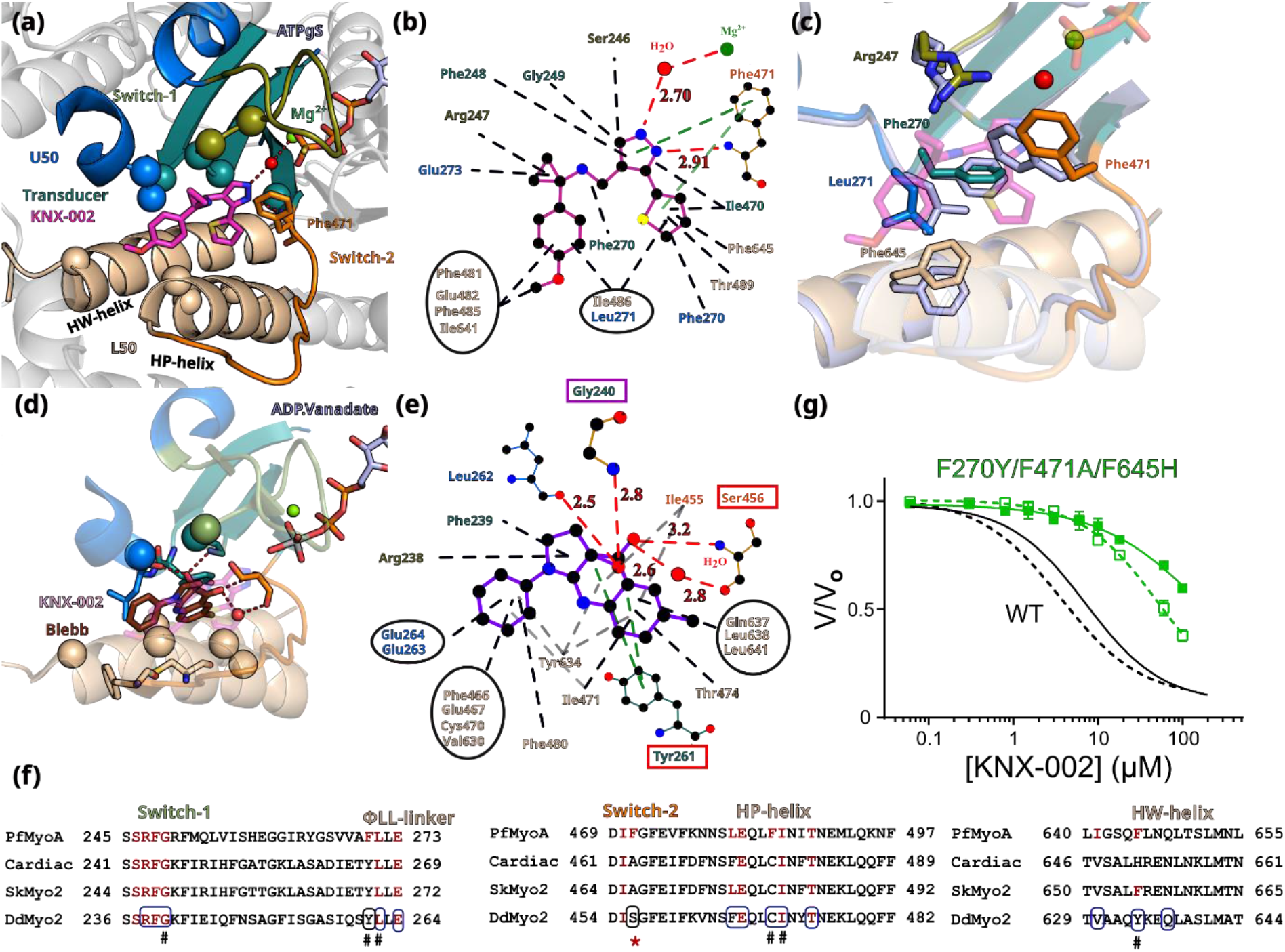
The inner pocket in which KNX-002 binds greatly differ from that of blebbistatin. **(a)** Zoom on the pocket of KNX-002. The regions involved in the binding are colored: Switch-2 (orange); deep teal cyan (Transducer); U50 (marine blue); L50 (wheat). Residues involved in binding are displayed as sphere, except for those involved in specific bonds (electrostatic or π-stacking). **(b)** Schematic representation of the interactions around KNX-002. Polar interactions are represented in red dash lines; π-stacking in green and apolar in black. **(c)** Superimposition of PfMyoA/KNX-002 (colored by subdomains) and PfMyoA/Apo (grey-blue). The residues with different conformation are shown as stick, indicating how adjustments are required to form the KNX-002 pocket. **(d)** Representation of the Blebbistatin (Blebb) binding pocket in Dictyostelium discoidum myosin 2 (DdMyo2, PDB code 1YV3,^27^ with orientation of the U50 subdomain as shown in 3a for PfMyoA. **(e)** Schematic representation of the interactions around Blebb. Residues that are different from PfMyoA but bind Blebb are highlighted in a red box; conserved residues involved in different bond type are in a purple box. **(f)** Sequence alignment of different myosins: PfMyoA, β-cardiac myosin 2 MYH7 (cardiac), skeletal muscle myosin 2 (SkMyo2) and DdMyo2 analyzing the conservation of residues involved in the binding of KNX-002 and Blebb. Residues involved in KNX-002 binding are in red, they are also colored in other myosins when conserved. Residues involved in Blebb are circled. Residues involved in electrostatic or stacking interactions are labelled with a red star. Remarkable positions involved in drug binding and specificity are labelled with a hash. **(g)** KNX-002 poorly inhibits the actin-activated (filled green squares, IC_50_>100 μM) and the basal ATPase activity (open green squares, 52 μM (95% CI, 31-149) of the triple mutant F270Y/F471A/F645H compared to its effect on WT PfMyoA (WT fits reproduced from Fig. 1a, actin-activated (solid line) and basal (dashed black line) ATPase activity). n=2 for each assay.

Binding of the KNX-002 molecule, comprised of four moieties (**Fig. 1b**), seems to be stabilized mainly by the hydrophobic effect: the methoxyphenyl **(D)** and the methylcyclopropamine **(C)** are sandwiched between apolar groups of residues from the Switch-1, β7 strand of the Transducer and the ɸLL linker, a highly conserved linker located after the _Transducer_β7 and on the U50 side (**Supplementary Fig. 3c**), as well as apolar residues from the ^L50^HP- and ^L50^HW-helices (**Fig. 3a, 3b, Supplementary Table 2**). The pyrazole **(B)** and thiophene moieties **(A)** are stacked between residues from the β5 and β6 strands of the Transducer, the ^L50^HW-helix with contribution of Switch-2. Amongst these bonds, the most remarkable are established by the pyrazole (B), as it is involved in **(i)** a polar interaction with a water molecule coordinating the Mg^2+^ ion that interacts directly with the nucleotide and **(ii)** π-stacking electrostatic interactions with _Switch-2_Phe471 (**Fig. 3a, 3b, Supplementary Table 3**). The tight fit of KNX-002 in this pocket is remarkable because all four cyclic entities make at least one contact within a radius of ~3.4 Å with atoms of the pocket.

In addition to the small main-chain shifts to adopt the size of the pocket around KNX-002 (**Supplementary Fig. 3a**), side chains of key-binding residues rotate to fit KNX-002 binding (**Fig. 3c**). This is specifically true for _Switch-2_Phe471, _ɸLL linker_Leu271 or _HW-helix_Phe645 (**Fig. 3c**).

It is important to describe how the myo2 inhibitor Blebbistatin (Blebb) and KNX-002 binding pockets differ because they involve similar PfMyoA structural elements. Indeed, the Blebb binding pocket also involves the Switches-1 and -2, the Transducer and ɸLL linker, as well as the HP- and HW-helices (**Fig. 3d**). However, while the binding of KNX-002 and Blebb partially involve the same residues, the conformation of the pockets are in fact very different and thus Blebb cannot fit in the KNX-002 pocket and vice-versa. The two compounds indeed target distinct structural ATP-states of the motor: KNX-002 binds the PR state while Blebb targets the PPS state ^27,16^. Moreover, Blebb and KNX-002 have different scaffolds and although binding of both mainly involves apolar interactions, the nature of the interactions and the residues involved differ (**Fig. 3b, 3e**). In addition, the nature of the few polar interactions are drastically different. For example, the hydroxyl moiety of Blebb is involved in essential electrostatic interactions with the amide of _Switch-1_Gly240, the carbonyl of _ɸLL Linker_Leu262 and π-stacking interactions are established between Blebb and the _ɸLL linker_Y261 (Dd, 1YV3, **Fig. 3e**), while only apolar interactions are made with the PfMyoA equivalent _Switch-1_Gly249, _ɸLL linker_Leu271 and _ɸLL linker_Phe270 residues and KNX-002. The Switch-2 conformation as found in the pre-powerstroke state excludes the possibility of a direct bond between blebbistatin and the nucleotide or the Mg^2+^ ion. In contrast, specific interactions occur for KNX-002 when the Switch-2 adopts the conformation found in the Post-rigor state. Taken together, these results clearly demonstrate that KNX-002 binds a novel and previously undescribed pocket with unique features. The most interesting novel characteristic of the KNX-002 pocket is that it involves the binding of a water that coordinates both the Mg^2+^ ion and the nucleotide.

Sequence polymorphism between Myo2s and PfMyoA explain the specificity of the two inhibitors for their corresponding binding sites and result in different specific requirement for their binding mode. In particular, KNX-002 specificity for PfMyoA results in particular from three major sequence change compared to Myo2s: **_Switch-2_Phe471 in PfMyoA**, involved in essential π-stacking interactions with KNX-002, is replaced by a serine or alanine in Myo2s (_Switch-2_Ser456 in DdMyo2 (1YV3)); **_HP-helix_Phe485 in PfMyoA** is replaced by a cysteine in DdMyo2 (_HP-helix_Cys470); **_HP-helix_Leu481 in PfMyoA** is replaced by a Phenylalanine (_HP-helix_Phe466 in DdMyo2) (**Fig. 3f**). The Phe/Leu polymorphism in the HP-helix was already reported as responsible of high specificity amongst Myo2s for a Blebb derivative of high therapeutical potential (MPH-220) ^16^. Such sequence differences allow each inhibitor to interact specifically in its tight pocket.

We validated this site and the role of some residues in KNX-002 binding and specificity with a triple mutant of PfMyoA. Three residues of the PfMyoA sequence were replaced by that of β-cardiac muscle myosin (F270Y/F471A/F645H) at key-positions of the KNX-002 binding (**Supplementary Fig. 4, Fig. 3f**). As in cardiac myosin (**Fig. 1a**), this triple mutant became less sensitive to KNX-002 compared with WT, with basal ATPase IC_50_ increasing from 3.6 to 52 μM, and actin-activated ATPase IC_50_ increasing from 7.2 to >100 μM (**Fig. 3g**). This result not only validates the binding pocket of KNX-002, but also suggests that some or all of these three key-aromatic positions are essential in the efficient binding and specificity of KNX-002 for PfMyoA.

### KNX-002 slows ATP binding and affects Mg^2+^ coordination in the active site

Compared to Blebb, KNX-002 is positioned closer to the active site and the water that binds strongly to its pyrazole group also participates in Mg^2+^ and γP_i_ coordination, suggesting that KNX-002 could influence ATP binding, in addition to its hydrolysis. To test the hypothesis of a direct effect of KNX-002 on ATP binding and gain insights into the mechanism of action of the compound, we used transient kinetics to measure the effect of KNX-002 on Mant-ATP and Mant-ADP binding to PfMyoA. KNX-002 reduces the affinity of PfMyoA for ATP 11-fold (Kd 0.29 μM vs. 3.2 μM, **Fig. 4a**), while the effect is more subtle for ADP (Kd 2.9 μM vs. 3.9 μM, **Supplementary Fig. 5**). Since Mg^2+^ interacts directly with the water coordinating KNX-002 and ATP, we also performed ATP binding experiments at lower Mg^2+^ concentration. As expected, the ability of KNX-002 to reduce ATP affinity was not as significant at low magnesium (Kd reduced 3.6-fold, **Fig. 4a**), consistent with the essential role of the water coordination to the Mg^2+^ ion in KNX-002 mode of action when myosin binds ATP.

**Figure 4.**
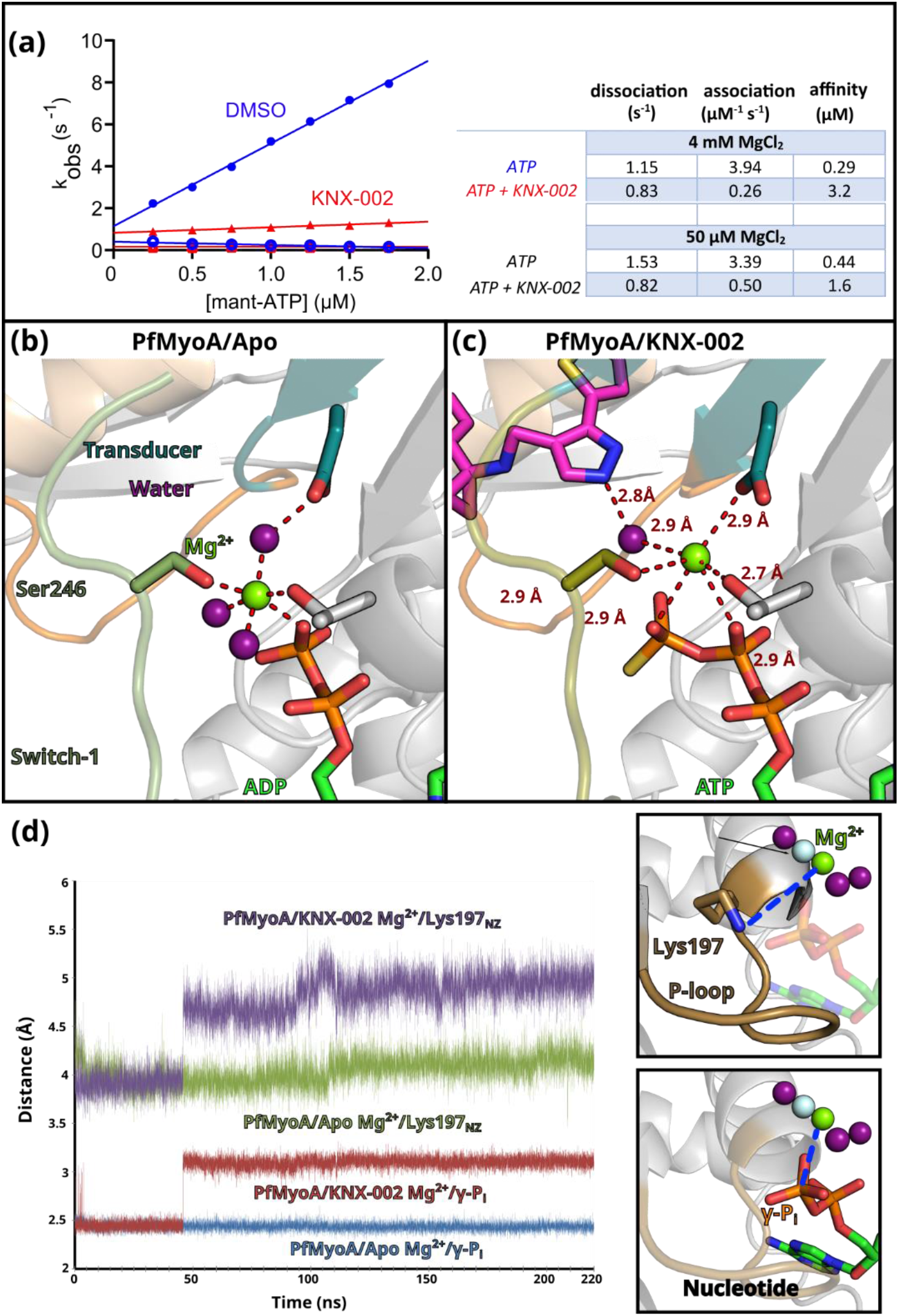
KNX-002 alters the coordination of the Mg^2+^ ion. **(a)** KNX-002 slows down mant-ATP binding and weakens the affinity of PfMyoA for ATP. Rates in the absence (blue circles) or presence of 100 μM KNX-002 (red triangles) of the fast (filled symbols) and slow phases (open symbols) from the biphasic transients at 4 mM MgCl_2_ are plotted. The association rate (slope) and dissociation rate (y-intercept) are given in the Table at high and low MgCl_2_ concentration. Data represent two experiments with independent protein preparations. (**b** and **c**) The difference in the Mg^2+^ ion coordination is illustrated by comparing structures obtained in the absence of KNX-002 (**b -** PfMyoA/Apo) or in the presence of KNX-002 (**c** - PfMyoA/KNX-002), see also Fig. 6. KNX-002 remodels the coordination of the Mg^2+^ ion through a direct interaction with a water involved in the coordination of both the Mg^2+^ ion and γP_i_. **(d)** Evolution of two reporters of the Mg^2+^ position and coordination: plotted are the distance between Mg^2+^ and the nitrogen of the side-chain of the catalytic Lysine 197 (Lys197_NZ_) and the distance between Mg^2+^ and the γ-phosphorus (γ-P_i_) of the ATP. **(Right)** The two reporters are shown on a cartoon representation, the green and purple spheres correspond to the Mg^2+^ position and three coordinating waters as found in the PfMyoA-Apo structure, the grey sphere indicates the Mg^2+^ position in the KNX-002-PfMyoA structure.

A detailed comparison of the apo and KNX-002-bound structures reveals that the interaction between KNX-002 and the water near the _KNX-002_pyrazole group directly alters the coordination of the Mg^2+^ ion (**Fig. 4b, 4c**). This change in the coordination of the Mg^2+^ also results in a slightly different position of the ion in presence of KNX-002 (**Fig. 4c**). Note in particular the longer distances around the Mg^2+^ and the fact that it interacts directly with the _Switch-2_Asp469 when KNX-002 is present, while for PfMyoA/Apo, a water molecule mediates this interaction and a perfect hexa-coordination is formed. The Mg^2+^ position and hexa-coordination is the same when either ATP or ADP are bound in the post-rigor state (**Supplementary Fig. 6**). The fact that the β-phosphate is further away from KNX-002 than the γ-phosphate explains why KNX-002 affects ADP binding to myosin to a lesser degree compared to ATP (**Fig. 4a**; **Supplementary Fig. 5**).

To further investigate how KNX-002 alters the coordination and the position of the Mg^2+^ ion, we performed all-atom molecular dynamics simulations during 220 ns with Mg.ATP bound in an apo and KNX-002 bound structures (see Methods). In the two simulations, the starting position of the Mg^2+^ ion was imposed as found in the apo structure. During the entire time course of the simulation, both the drug and the nucleotide stayed in place. To evaluate how the presence of KNX-002 affects it, two reporter distances were chosen to monitor the relative position of the Mg^2+^ ion **(i)** the distance with the nitrogen of the P-loop Lys197 which is also in close vicinity of the γP_i_ and waters from the Mg^2+^ coordination and **(ii)** the distance with the γ-phosphorus (γ-P). These two distances were plotted during time (**Fig. 4d**). While in the Apo condition, the distances of Mg^2+^ with both reporters are stable, they rapidly increase around 40 ns when KNX-002 is present. The distances monitor a deviation in the position of the Mg^2+^ and its coordination, confirming the direct effect of KNX-002 on the Mg^2+^ ion.

Taken together, these results demonstrate that PfMyoA adopts a stable **PR state when both ATP and KNX-002 are bound, in which the inner cleft cannot close, thus preventing the recovery stroke**. The water near the pyrazole undergoes a trade-off to bind either KNX-002 or the Mg^2+^ ion. This destabilizes the coordination around the Mg^2+^ ion. Given the essentiality of Mg^2+^ ion in both ATP binding, hydrolysis and product release in myosins ^28,29,30^, direct alteration of the Mg^2+^ coordination by KNX-002 is a novel and key mode of action for this class of myosin inhibitors.

### PfMyoA.ADP has low affinity for F-actin in the presence of KNX-002

To investigate whether KNX-002 affects the stability of PfMyoA actin-bound states, we performed actin pull-down experiments using surface-bound PfMyoA (see Methods, **Fig. 5a, Supplementary Fig. 7, Supplementary Fig. 8**) in the presence of either ATP, ADP, or no nucleotide with or without KNX-002.

**Figure 5.**
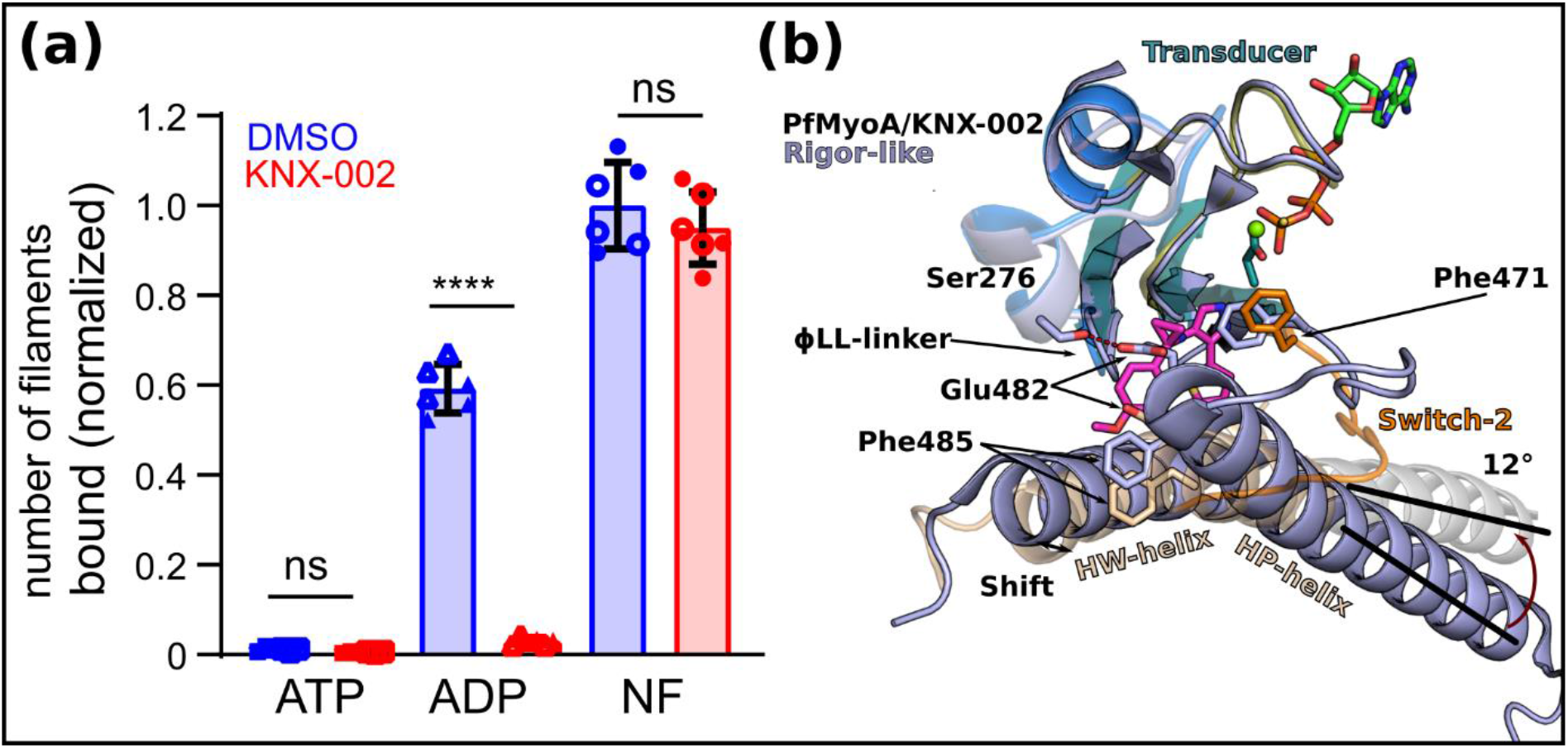
PfMyoA.ADP binds actin weakly in the presence of KNX-002. **(a)** KNX-002 weakens the affinity of actin for M.ADP but has no effect on binding in the presence of ATP or with nucleotide-free (NF)-PfMyoA. Number of actin filaments bound ± SD to surface immobilized PfMyoA (see Methods). See **Supplementary Fig. 7** for examples of the raw data and **Supplementary Fig. 8** for data with an expanded y-axis scale. The difference between ADP ± KNX-002 is significant (p<0.0001) (ANOVA followed by Tukey’s post-hoc test), but there were no significant differences between ATP ± KNX-002 (p>0.999), nor between NF ± KNX-002 (p=0.64). Data represent two experiments (open and filled symbols) using independent protein preparations. **(b)** The KNX-002 binding pocket does not exist in the Rigor state. PfMyoA/KNX-002 (colours) and PfMyoA in the Rigor-like state (light blue) (PDB code 6I7D, ^9^ Robert-Paganin et al., 2019) are superimposed on the U50 subdomain and show how a change in the conformation and the orientation of the L50 subdomain would close the inner cleft and would thus not be compatible with KNX-002 binding. Indeed, the ^L50^HP-helix rotates by 12° and the ^L50^HW-helix shifts, changing the position of F485 and F645, two residues involved in KNX-002 binding. The reorientation of Switch-2 also leads to a new position for the key F471 residue that is incompatible with KNX-002 docking into this site in the Rigor state.

As expected, myosin binds few filaments in the presence of ATP, and KNX-002 does not change this property (**Fig. 5a**). In the absence of nucleotide (Nucleotide Free, NF), a similar large number of actin filaments are recruited in both Apo and KNX-002 conditions. The absence of nucleotide allows myosin to bind with high affinity for actin in the Rigor state, a state with the actin-binding cleft closed ^31^(**Supplementary Fig. 2**). Note that in this Rigor state, the ɸLL linker directly interacts with the HP and HW helices and thus the KNX-002 binding site we identified does not exist. The observation that KNX-002 does not change the amount of PfMyoA bound to actin in NF conditions is consistent with the fact that PfMyoA stays bound in the Rigor state with the inner pocket closed, which would prevent KNX-002 binding (**Fig. 5b**). KNX-002 also does not affect the rate of ATP induced actomyosin dissociation, further suggesting that the compound is not compatible with the Rigor state (**Supplementary Fig. 9**). In ADP, fewer actin filaments are recruited compared to the NF condition. More importantly, the presence of both ADP and KNX-002 leads to a level of actin binding comparable to the weak binding observed with ATP (**Fig. 5a**). This result suggests that PfMyoA.ADP binds KNX-002 and prevents it from binding strongly to the F-actin track. Taken together, these results suggest that KNX-002 favors trapping the motor in a state of poor affinity for the actin track when nucleotide is bound.

While we crystallized the motor in the PR state in the presence of ATPγS, crystals were also obtained with MgADP in the same Post-rigor state. Since A.M.ADP complexes are in equilibrium with detached M.ADP motors, it is likely that KNX-002 can bind these detached M.ADP motors as they adopt the post-rigor state. KNX-002 binding would thus stabilize this state and prevent re-attachment to the actin filament in a Strong-ADP bound state. The current structures available for distinct myosins in the Strong-ADP bound state ^25^ have indeed shown that the inner cleft is closed in these structures. Thus, KNX-002 binding in the inner cleft would prevent strong binding to the track (**Fig. 5b**).

## Discussion

KNX-002 is a first-in-class inhibitor of PfMyoA. As previously shown from knockout experiments, PfMyoA is an essential myosin for *Plasmodium falciparum* invasion and motility, and as such represents a novel and previously untapped target that could be used to prevent *Plasmodium falciparum* pathogenesis ^9,10^. The fact that KNX-002 ablates *P. falciparum* blood stage growth highlights the potential of targeting PfMyoA to develop future antimalarial drugs.

Previous studies of myosin inhibitors have shown that these compounds target binding pockets that are not always present in the apo form ^17^ and that the accessibility of these cryptic pockets may vary depending on the myosin classes and the structural states targeted ^17^. This is why the binding mode of most myosin modulators cannot be predicted. Here we show that KNX-002 binds in the inner pocket of PfMyoA via structural and biochemical data and validated this site via directed mutagenesis. While KNX-002 and Blebb binding involves the same structural elements: i.e. Switch-1 and Switch-2, HP-helix, HW-helix and the ɸLL-linker, the conformation of the pocket strongly differs. This is due to the sequence polymorphism between myosins-2 and PfMyoA, as well as the fact that the two compounds target different structural states. Unlike previous inhibitors binding via residues found in the inner pocket, such as the Pre-power stroke state targeted by Myo2 inhibitors such as Blebb and its derivatives, KNX-002 binds to a myosin conformation called the Post-rigor state. KNX-002 thus targets a previously undescribed binding pocket, namely the inner pocket of a Post-rigor myosin state. Based on sequence alignments, this binding pocket appears to be conserved in MyoA from other apicomplexan parasites such as *Toxoplasma* (toxoplasmosis), *Eimeria* (coccidiosis), and *Babesia* (babesiosis) and thus derivatives of KNX-002 could ultimately be applicable to treatment of a number of parasitic diseases (**Supplementary data 1**).

Interestingly, transient kinetics and structural data both demonstrate that the mechanism of action of these inhibitors greatly differ. While Blebb inhibition comes from the occupation of the inner pocket and blocking efficient initiation of the powerstroke, it does not interfere with ATP hydrolysis^32,27,16^. In contrast, KNX-002 steric occupation of the inner cleft in the post-rigor state pocket decreases ATP hydrolysis. Our structure suggests that the molecular basis for reduced ATP hydrolysis is the disruption of Mg^2+^ coordination via interaction of KNX-002 with a water molecule, thus interfering with nucleotide binding (**Fig. 6**). Blebb and KNX-002 therefore belong to different categories of inhibitors with different mechanisms of action. As the first representative of a novel class of inhibitors, the mode of action of KNX-002 results from sequestering an “incompetent” post-rigor state. KNX-002 indeed stabilizes a state of low affinity for actin in which binding to the inner cleft prevents the PfMyoA recovery stroke and in which disruption of Mg^2+^ coordination via a water mediated direct interaction with KNX-002 contributes to preventing efficient ATP hydrolysis. Importantly, it is the first time that a compound is reported to bind to the inner pocket of a Post-rigor myosin state, without the requirement for much conformational change. This opens the door for the potential discovery of distinct inhibitors for different myosin targets as we report here an efficient and novel way of preventing myosin from participating in force production.

**Figure 6.**
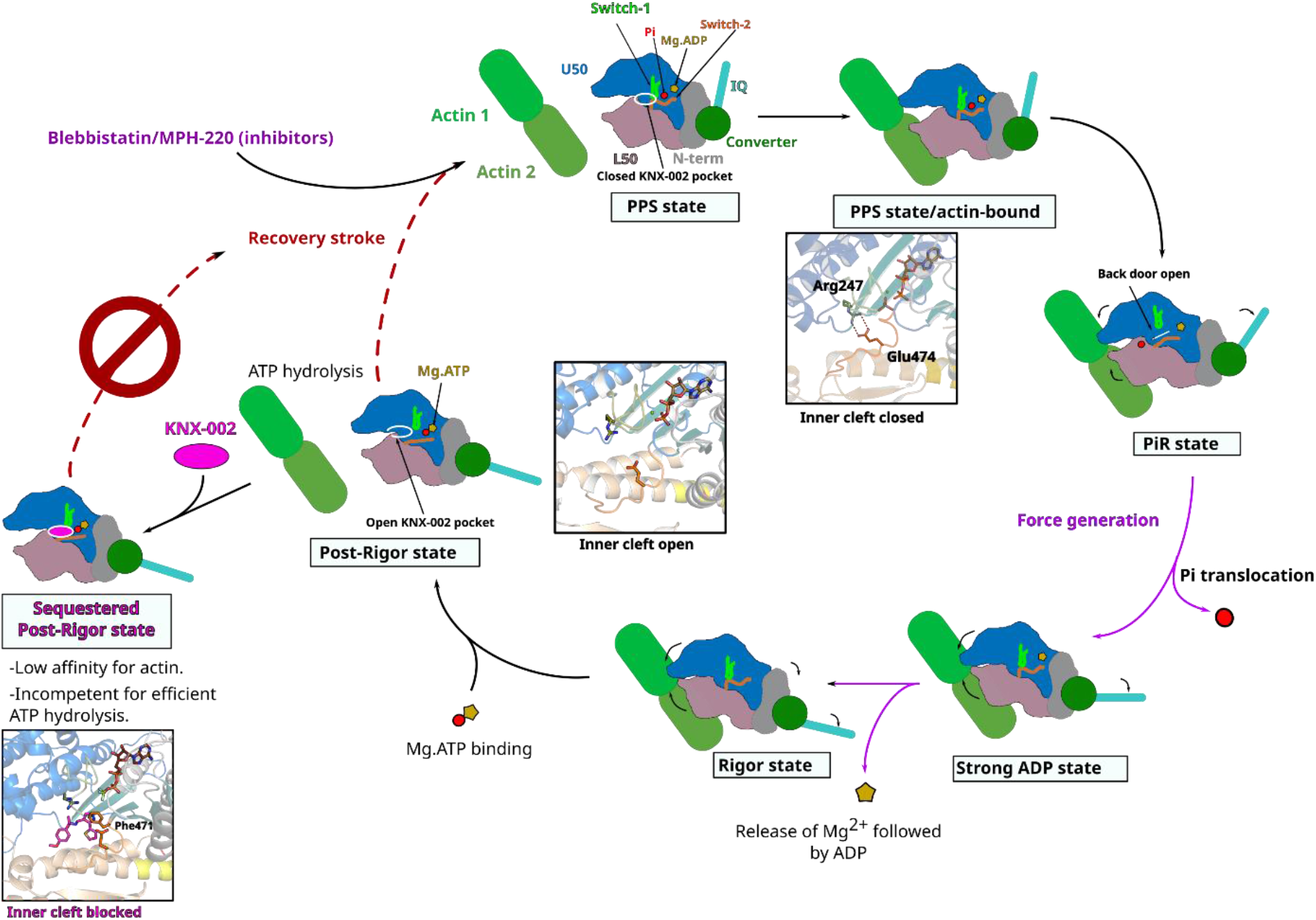
Mechanism of the inhibition by compound KNX-002. Schematic representation of the motor cycle of PfMyoA, where the state of nucleotide bound governs **(i)** the conformation of the motor and **(ii)** the affinity for the actin filament. In the post-rigor (PR) state, PfMyoA binds Mg.ATP and has low affinity for actin since the actin-binding cleft between U50 and L50 subdomains is open. After the recovery stroke that primes the lever arm, PfMyoA adopts the pre-powerstroke (PPS) state in which ATP is hydrolyzed. The weak association of the PPS to actin initiates a transition towards the Pi release state (PiR), the first force-producing state. This initiation of the powerstroke allows the opening of the Pi release tunnel allowing Pi translocation. After Pi release, a large swing of the lever arm (powerstroke) leads to a Strong ADP state in which the actin-binding cleft is closed allowing stronger association with actin. Mg^2+^ and ADP are finally released from the Rigor state after a small swing of the lever arm and reorientation of the N-terminal subdomain. An ATP molecule can bind and leads to a fast transition towards the Post-rigor state that detaches from actin. In contrast to the previously described Myo2 inhibitors, Blebbistatin and MPH-220 ^35^ which target PPS, KNX-002 targets PR, which prevents the recovery stroke and ATP hydrolysis. When KNX-002 intercalates between the U50 and the L50, it stabilizes a state of poor affinity for F-actin that is not compatible with efficient ATP hydrolysis.

The ability of KNX-002 to inhibit asexual blood stage growth *in vitro* demonstrates its antimalarial activity. It confirms that PfMyoA and myosins in general are potential antiparasitic pharmacological targets ^9,10,31,23^. While the IC_50_ for KNX-002 is in the micromolar range, these current data set a foundation for substantial further development with the structure of the PfMyoA bound to KNX-002 able to direct SAR that promote higher potency while conserving specificity. Thus the structural data we provide opens new roads to optimize this molecule for therapeutical potential. The biggest advantage of a strategy targeting PfMyoA with a KNX-002 derivative is that by inhibiting actin binding, it should efficiently slow motility and invasion phases of the parasite. Future studies will aim to bring new prophylactic and/or symptomatic treatments. In addition, it will be essential to study how such potent inhibitors could bring a significant advantage either in conjunction with other antimalarials ^33,34^ or in conjunction with vaccination strategies that are unfortunately poorly efficient at present ^6^.

## Abbreviations

Blebb: blebbistatin;
NF: Nucleotide-free
Mant-ATP: 2’/3’-O-(N-Methyl-anthraniloyl)-adenosine-5’-triphosphate

## Acknowledgements

This work was supported by NIH grant R01AI 132378 to KMT and AH, a grant from the Human Frontier Science Program (RGY006/2016) to JB and AH, grants from the Wellcome (Investigator Award to JB, 100993/Z/13/Z) and the Medicines for Malaria Venture (RD-08-2800) to JB. We thank Cytokinetics for providing purified actin and access to their high-throughput screening library, infrastructure and staff. We are grateful to Dr. James Spudich (CEO of Kainomyx) for giving us the opportunity to study KNX-002 in the Plasmodium system via a Continuing Research Agreement between Kainomyx and the University of Vermont (KMT), and Kainomyx and the Institut Curie (AH). We acknowledge the ESRF for providing in-house beamtime on the ID30B. We thank Dr. James Spudich and Dr. Kathleen Ruppel for critical reading of the manuscript, and James Spudich for useful suggestions regarding the motility experiments. We thank Dr. Gordon Leonard (ESRF, Grenoble, France) for critical reading of the manuscript.

## Competing interests

A.H. receives research funding from Cytokinetics and consults for Kainomyx, inc.

## Materials and methods

### KNX-002

KNX-002 (formerly CK2140597) was originally identified from high throughput screening of a library of 50000 compounds at 40uM final concentration (2% DMSO, v/v), performed by JS and EW at Cytokinetics, to identify inhibitors of the actin-activated ATPase of *T. gondii* MyoA (protein provided by Gary Ward, University of Vermont), and PfMyoA (protein provided by KMT). A coupled enzymatic ATPase assay^9^ was used for screening. Actin was first polymerized using 1 mM ATP, 2 mM MgCl_2_, 20 mM KCl and left on ice for 30 min. Solution A was made with PM12 (12 mM Pipes-KOH, pH 7.0, 2 mM MgCl_2_), 1 mM DTT, 0.1 mg/ml BSA, 0.4 mM pyruvate kinase/lactate dehydrogenase, 1 mg/ml Myosin XIV, 0.009% Anitfoam (Sigma-Aldrich). Solution B was made with PM12, 1 mM DTT, 0.1 mg/ml BSA, 1 mM ATP, 1 mM NADH, 1.5 mM phosphoenolpyruvate, 0.6 mg/ml actin previously polymerized and 0.009% Antifoam. 12.5 μl each of Solution A and B were added to a plate with a Packard MiniTrak liquid handler and then mixed at 2400 rpm for 45 s with an BioShake 3000 ELM orbital plate mixer (Qinstruments). The decrease in optical density at 340 nm as a function of time was measured on an Envision 2104 spectrophotometer (Perkin Elmer) with a 600 s data monitoring window sampled every 60 s. Hit progression included confirmation of activity in an independent experiment, inactivity against an unrelated ATPase (hexokinase), and dose-responsive inhibition. Hexokinase reactions were performed using glucose (1 mM) and sufficient hexokinase to generate ADP from ATP at a rate similar to the target myosins. Hit progression included confirmation of activity in an independent experiment, inactivity against an unrelated ATPase, and dose-responsive inhibition. Of note, KNX-002 was initially a weak hit in both campaigns due to a technical issue, but pursued for TgMyoA and later confirmed to be robustly active in both TgMyoA and PfMyoA assays, both with the initial library batch as well as with resupplied compound (>97% pure by LC/MS, Asinex). Kainomyx (www.kainomyx.com) licensed 6 compounds from Cytokinetics for drug discovery for malaria and other parasitic diseases, and James Spudich, CEO of Kainomyx, released CK2140597 (now KNX-002) without restrictions for academic studies in both the laboratories of Gary Ward, University of Vermont (for *Toxoplasma*), and KMT, University of Vermont (for *Plasmodium*). The KNX-002 used in this manuscript was custom-synthesized by Asinex (Winston-Salem, NC), and showed a high level of purity (97% pure), based on peak areas from a UV HPLC chromatogram at 275 nm using a Diode Array Detector.

### Myosin expression and purification

The full length PfMyoA heavy chain WT and mutants were co-expressed with PUNC chaperones and two lights chains PfELC and PfMTIP. FLAG-tagged recombinant proteins were purified from baculovirus-infected SF9 cell using previously described methods in ^36^. The constructs were purified for the crystallization assays. PfMyoA full length co-expressed with PfELC and PfMTIP with the 60 amino acids in N-term removed (PfMTIP-∆N) co-expressed with chaperones as described in ^10^. The cells were grown for 72 h in medium containing 0.2 mg/ml biotin, harvested and lysed by sonication in 10 mM imidazole, pH 7.4, 0.2 M NaCl, 1 mM EGTA, 5 mM MgCl2, 7% (w/v) sucrose, 2 mM DTT, 0.5 mM 4-(2-aminoethyl) benzene-sulfonyl fluoride, 5 μg/ml leupeptin, 2 mM MgATP. An additional 2 mM MgATP was added prior to a clarifying spin at 200,000 ×g for 40 min. The supernatant was purified using FLAG-affinity chromatography (Sigma). The column was washed with 10 mM imidazole pH 7.4, 0.2 M NaCl, and 1 mM EGTA and the myosin eluted from the column using the same buffer plus 0.1 mg/ml FLAG peptide. The fractions containing myosin were pooled and concentrated using an Amicon centrifugal filter device (Millipore), and dialyzed overnight against 10 mM imidazole, pH 7.4, 0.2 M NaCl, 1 mM EGTA, 55% (v/v) glycerol, 1 mM DTT, and 1 μg/ml leupeptin and stored at −20 °C. Skeletal muscle actin was purified from chicken skeletal muscle tissue essentially as described in ^37^.

### Basal and actin-activated ATPase assays

ATPase activity was determined using linked assay, which couples the regeneration of hydrolyzed ATP to the oxidation of NADH to NAD^+^ ^9^. The decrease in optical density at 340 nm as a function of time was measured on a Lambda 25 UV/VIS spectrophotometer (Perkin Elmer) with a 360 s data monitoring window sampled every 5 s. Data were fit with a four-parameter logistic curve (Graph Pad Prism). Conditions: 10 mM imidazole pH 7.5, 5 mM KCl, 1 mM MgCl_2_, 1 mM EGTA, 1 mM DTT, 1mM NaN_3_, 2 mM MgATP, 5 nM PfMyoA, 55 nM skeletal myosin II, 100 nM cardiac myosin, 50 μM skeletal actin, 1% DMSO, 30°C. Under these conditions, the actin-activated values for PfMyoA in the absence of inhibitor were 84.2 ± 1.4 s^−1^ (n=3) and 9.6 ± 3.4 s^−1^ in the presence of 100 μM KNX-002.

### Motility and actin binding assays

The *in vitro* motility assay was performed as described^9^, except that the methylcellulose concentration was reduced to 0.15%. Actin binding to myosin was measured using a modified *in vitro* motility measurement method. All steps up to the triple rinse following neutravidin addition to the coverslip are the same as described in ^9^. From the step of myosin addition through to visualization, either 1% DMSO or 100 μM KNX-002 in 1% DMSO is kept in all solutions. For visualization in apo, ADP and ATP buffers, our motility buffer (buffer A, ^9^) that contained oxygen scavengers and 2 mM MgADP or 2 mM MgATP but no methylcellulose was added after the rhodamine-phalloidin actin binding step. The number of filaments bound was quantified by dividing each picture of actin filaments into four equal quadrants. One quadrant was picked at random and all filaments within the quadrant were counted by hand. The quadrant selected for quantification was held constant for all conditions. Conditions: 25 mM imidazole pH 7.5, 150 mM KCl, 4 mM MgCl_2_, 1 mM EGTA, 10 mM DTT, 0% methylcellulose, 1% DMSO, 30°C. 2 mM MgATP or MgADP were added for +ATP or +ADP condition, respectively.

### Phosphate burst

Chemical hydrolysis of ATP by myosin was performed by manual mixing under single turnover conditions. PfMyoA (25 μM) was pre incubated with 1% DMSO or KNX-002 (100 μM in 1% DMSO), manually mixed with 20 μM ATP, aged for 5 s, and then quenched by addition of 0.3 M perchloric acid before quantifying free phosphate using malachite green (Fisher), ^38^. Conditions: 10 mM imidazole pH 7.5, 150 mM KCl, 4 mM MgCl_2_, 1 mM EGTA, 10 mM DTT, 25 μM PfMyoA, 20 μM MgATP, 1% DMSO, 30°C.

### Transient kinetics

All stopped-flow experiments were performed on a KinTek stopped-flow apparatus (KinTek Corporation, model SF-2001). All concentrations stated are after mixing into the stopped-flow cell, except where stated. Light scattering at 295 nm with a 295 nm interference filter orthogonal to the incident light was used to monitor the dissociation of actomyosin (0.2 μM actin and 0.15 μM PfMyoA). Mant-ATP and mant-ADP binding to myosin was measured by exciting at 290 nm and monitoring emission light that passed through a 400 nm long pass filter. All traces were analyzed using the software package provided by KinTek and fit to either single or double exponentials. Multiple time courses (usually 3-8) were averaged prior to curve fitting. Conditions: 10 mM HEPES pH 7.5, 50 mM KCl, 4 mM MgCl_2_, 1 mM EGTA, 1 mM DTT, 1% DMSO, 30°C.

### Crystallization, data processing, structure determination and refinement

The construct PfMyoA FL complexed with its light chains PfELC and MTIP-∆N was co-crystallized with and without KNX-002. PfMyoA/ELC/ MTIP-∆N at 10.6 mg/ml was incubated with 2mM Mg.ADP-gamma-S (ATPγS) for 20 minutes (Apo-PfMyoA). PfMyoA-KNX-002 crystals were obtained with the incubation of 2mM Mg.ADP-gamma-S (ATPγS) for 20 minutes and 0.5 mM KNX-002 for 40 minutes. All the samples were centrifugated at 11,000 x g for 15 minutes at 4°C before crystallizing assays. Both crystals of PfMyoA/ELC/MTIP-∆N were obtained at 4°C by the hanging drop vapor diffusion method from a 1:1 (v:v) of protein and mother liquor. The crystallization buffer is 2.0 M ammonium sulfate, 0.1M sodium HEPES pH 7,5, 4% PEG400. The crystals were transferred in the cryoprotection containing the mother liquor + 25% glycerol (v:v) and immediately flash frozen in liquid nitrogen.

X-ray datasets were collected at the ESRF Synchrotron, on the ID30B beamline ^39^, with the following parameters at 100 K: Apo-PfMyoA (λ=0.9677 Å, 46.8% transmission, flux start 7.54.10^11^ ph/sec), PfMyoA-Knx002 (λ=0.9687 Å, 1.2% transmission, flux start 9.89.10^12^ ph/sec). Diffraction datasets were processed using XDS ^40^ and AutoPROC ^41^. Both crystal forms (Apo-PfMyoA) and (PfMyoA-KNX-002) belong to the space group P2_1_2_1_2_1_. The molecular replacement solution was obtained by using Phaser ^42^ using the PDB coordinates from 6YCY ^10^ without solvent and water as model target. The 3D structure of compound KNX-002 and the related restraints were generated using Elbow ^43^. The models were manually refined using Coot ^44^. The refinement was performed using Buster ^45^ and Phenix ^46^. The final Ramachandran for favored, allowed and outliers are 97.25%, 2.65%, 0.09% for (PfMyoA/Apo) and 97.08%; 2.92%, 0% for (PfMyoA/KNX-002), respectively. Coordinates of both structures are deposited in the PDB ^47^ under 8A12 and 8A13.

### Molecular dynamics simulations

Starting from the PfMyoA/Apo and PfMyoA/KNX-002 coordinates, ATP was modelled after ATPγS and the Mg^2+^ was positioned as found in PfMyoA/Apo. All molecular dynamics simulations were performed with Gromacs 2018.3 ^48^ on all-atom systems parametrized with charmm36m forcefield and built with CHARMM-GUI server ^49^. All systems consisted of a box with explicit water (TIP3) and neutralized with salt (KCl reaching 150 mM). Long-range electrostatic interactions were handled using the particle mesh Ewald (PME) method ^50^. The simulations were performed in an NPT ensemble; the temperature and the pressure of the system were fixed at 310.15 K with the Nosé-Hoover thermostat and 1 bar with the Parrinello-Rahman barostat. Trajectories of 220 ns were generated and further analyzed with the Gromacs tools and visualized in PyMOL ^51^ which served to create the illustrations.

### Effects of KNX-002 on parasite asexual blood-stage growth

Asexual parasite blood stage growth (measuring the ability of parasites to invade, replicate, exit and reinvade erythrocytes) was undertaken using the SYBR Green asexual growth assay, undertaken largely as published ^52^. In brief 96-well black clear bottom plates (Corning) were pre-printed with test compound and normalised with DMSO (Merck) to 0.5% of a total assay volume of 100μl. Highly synchronised ring stage parasites (see section 2.2.4) and blood was added to each well so that the final parasitaemia was 2% and the haematocrit was 1%. The compound was incubated with parasites for 72hrs before being frozen at −20°C (to aid with cell lysis). The plate was thawed on ice and a lysis buffer (20mM Tris pH7.5 (Merck), 5mM EDTA (Merck), 0.008% w/vol saponin (Merck), 0.08% vol/vol triton-x100 (Merck) and SYBR-Green I (Thermo Fisher Scientific) at a final concentration of 0.02% vol/vol). The 96-well plate was incubated for 1hr at RTP before each well was assayed for fluorescence using GFP filters (Excitation 485 nm/Emission 535 nm) on a microplate reader (TECAN).

### Toxicity assays

Cell lines were obtained from the American Type Culture Collection (Rockville, MD, USA) and were cultured according to the supplier’s instructions. Human MRC-5 *cell* line derived from normal lung tissue and HepG2 cell line derived from a hepatocellular carcinoma were grown in DMEM supplemented with 10% fetal calf serum (FCS) and 1% glutamine. Human hTERT-RPE1 cells were cultured in DMEM/F12 medium containing 10% fetal calf serum and 1% glutamine. Cells were maintained at 37°C in a humidified atmosphere containing 5% CO_2_.

Cell viability was determined by a luminescent assay according to the manufacturer’s instructions (Promega, Madison, WI, USA). Briefly, the cells were seeded in 96-well plates (2.5 × 10^3^ cells/well) containing 90 *μ*L of growth medium. After 24 h of culture, the cells were treated with the tested compounds at 10, 20, 50, 100 and 200 μM final concentrations. Control cells were treated with the vehicle.

After 72 h of incubation, 100 *μ*L of CellTiter Glo Reagent was added for 15 min before recording luminescence with a spectrophotometric plate reader PolarStar Omega (BMG LabTech). The percent viability index was calculated from three experiments.

## Supplementary data

**Supplementary Figure 1.**
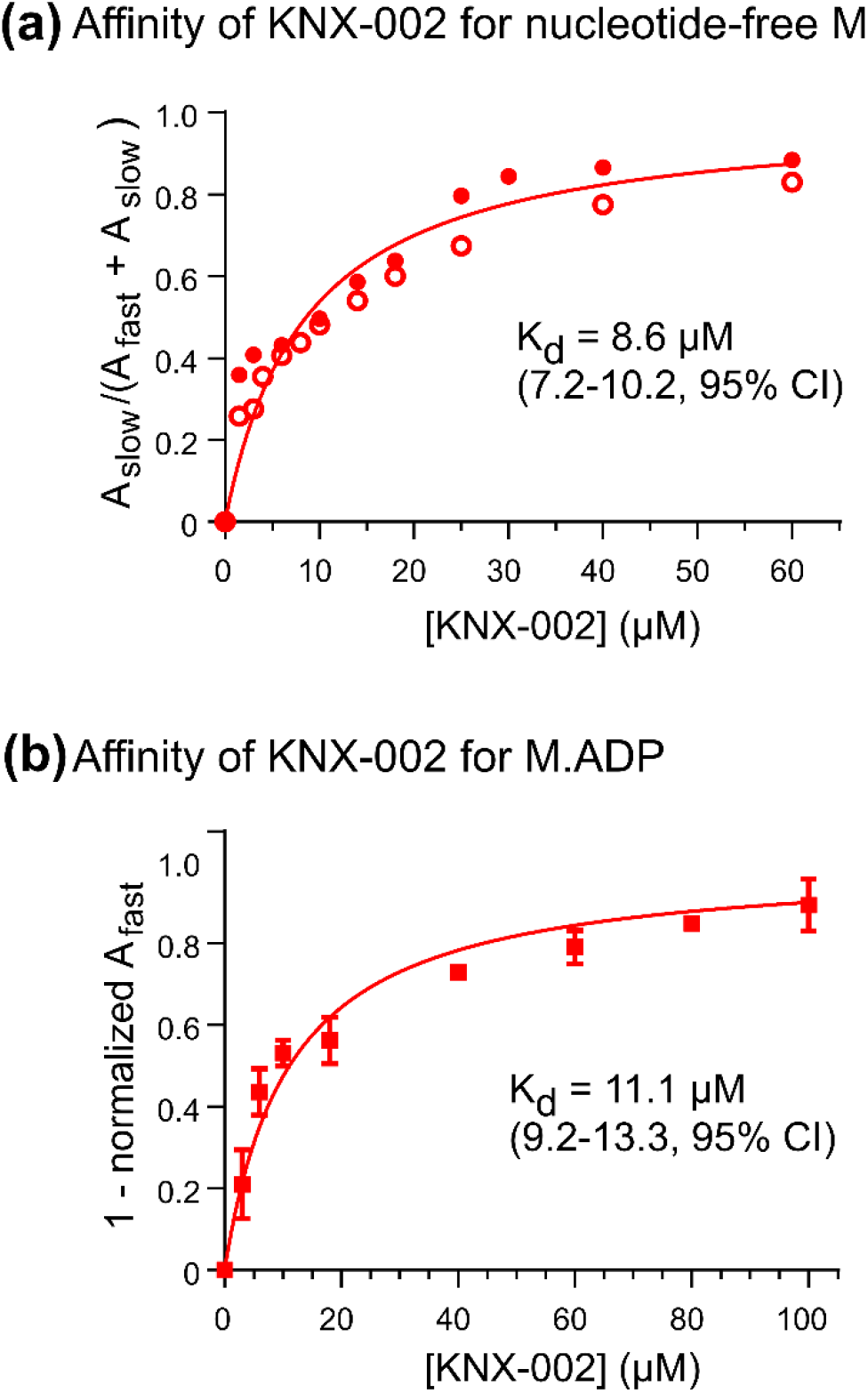
Affinity of KNX-002 for nucleotide-free PfMyoA and PfMyoA.ADP. **(a)** Affinity of KNX-002 for nucleotide-free M. Fractional amplitude of the slow phase (A_slow_/(A_fast_ + A_slow_)) as a function of KNX-002 concentration. 0.3 μM PfMyoA was pre-incubated with the indicated amount of KNX-002 and then rapidly mixed with either 3 μM mant-ATP (filled circles) or 6 μM mant-ADP (open circles) also containing the indicated amount of KNX-002. For mantATP, observed rate constants for the time courses were 6.5 ± 0.4 s^−1^ (no KNX-002) and 1.03 ± 0.01 s^−1^ (100 μM KNX-002). For mantADP, observed rate constants for the time courses were 12.7 ± 0.2 s^−1^ (no KNX-002) and 1.80 ± 0.02 s^−1^ (100 μM KNX-002). For intermediate KNX-002 concentrations, the fractional amplitude of the slow phase (A_slow_/(A_fast_ + A_slow_)) of the biphasic transients is plotted against KNX-002 concentration. From a hyperbolic fit to the data, the K_d_ of KNX-002 for nucleotide-free PfMyoA is 8.6 μM (7.2 – 10.2 μM). Value (95% CI). Two independent experiments were performed. Conditions: 10 mM HEPES pH 7.5, 50 mM KCl, 4 mM MgCl_2_, 1 mM EGTA, 1 mM DTT, 1% DMSO, 30°C. **(b)** Affinity of KNX-002 M.ADP. The disappearance of the fast phase (1-(A_fast_ at measured [KNX-002] / A_fast_ in the absence of KNX-002)) as a function of KNX-002 concentration. 0.3 μM PfMyoA was pre-incubated with 100 μM ADP and the indicated amount of KNX-002 and then rapidly mixed with 3 μM mant-ATP also containing the indicated amount of KNX-002. A hyperbolic fit to the data yields a K_d_ of KNX-002 PfMyoA.ADP of 11.1 μM (9.2 – 13.3 μM). Value (95% CI). All time courses were biphasic with a fast phase rate of 1.44 ± 0.03 s^−1^ of varying amplitude and a slow phase having a constant rate (0.08 ± 0.01 s^−1^) and amplitude. Two independent experiments were performed. Conditions: 10 mM HEPES pH 7.5, 50 mM KCl, 4 mM MgCl_2_, 1 mM EGTA, 1 mM DTT, 1% DMSO, 30°C.

**Supplementary Figure 2.**
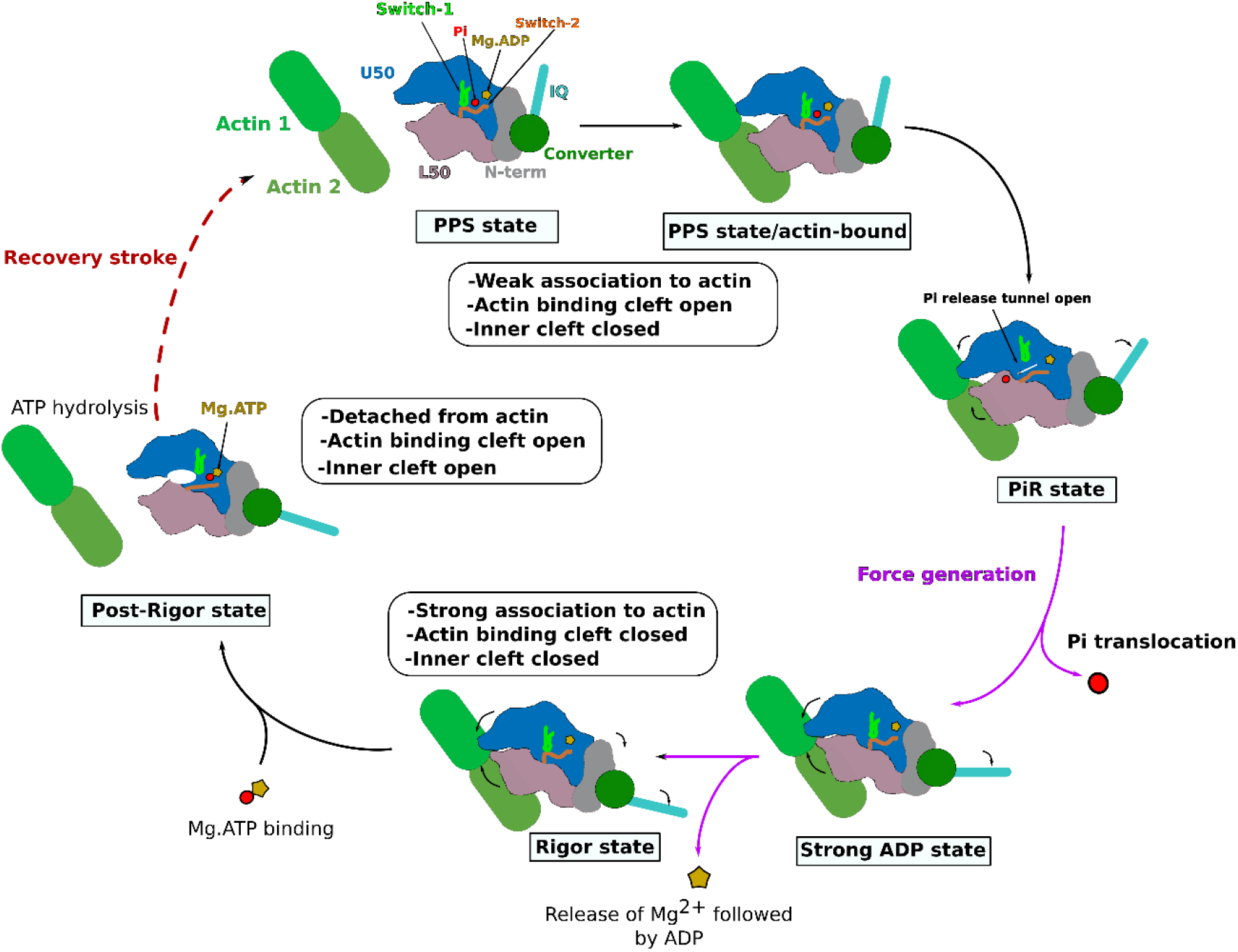
Schematic motor cycle of a myosin. In the post-rigor (PR) state, the myosin is bound to ATP, with the lever arm down and the actin binding cleft open, including the inner cleft. The PR is detached from actin. Priming of the lever arm occurs during the recovery stroke, resulting in the pre-power stroke state (PPS) in which ATP hydrolysis is facilitated thanks to the specific positioning of Switch-2 in this state (inner cleft or back-door closed). The PPS state can associate weakly to the actin track. This association triggers the transition towards the P_i_ release state (P_i_R) that allows P_i_ release. It is presumed that a small movement of the lever arm is sufficient to rearrange the actin-binding site in the P_i_R state so that an opening of the P_i_ release tunnel would then allow the P_i_ to escape the active site ^1^. The P_i_R state initiates force production. A large lever arm swing and cleft closure drive the transition to the Strong ADP state. Finally, the release of the Mg^2+^ ion followed by ADP occurs after the transition to the Rigor state. The Rigor state differs from the Strong ADP state by a further small lever arm swing. In the Rigor state, myosin is ready to bind a new ATP molecule. This leads to fast detachment from actin while myosin adopts a post-rigor state and starts a new cycle. The structural features underlying myosin force production are reviewed in ^2^.

**Supplementary Figure 3.**
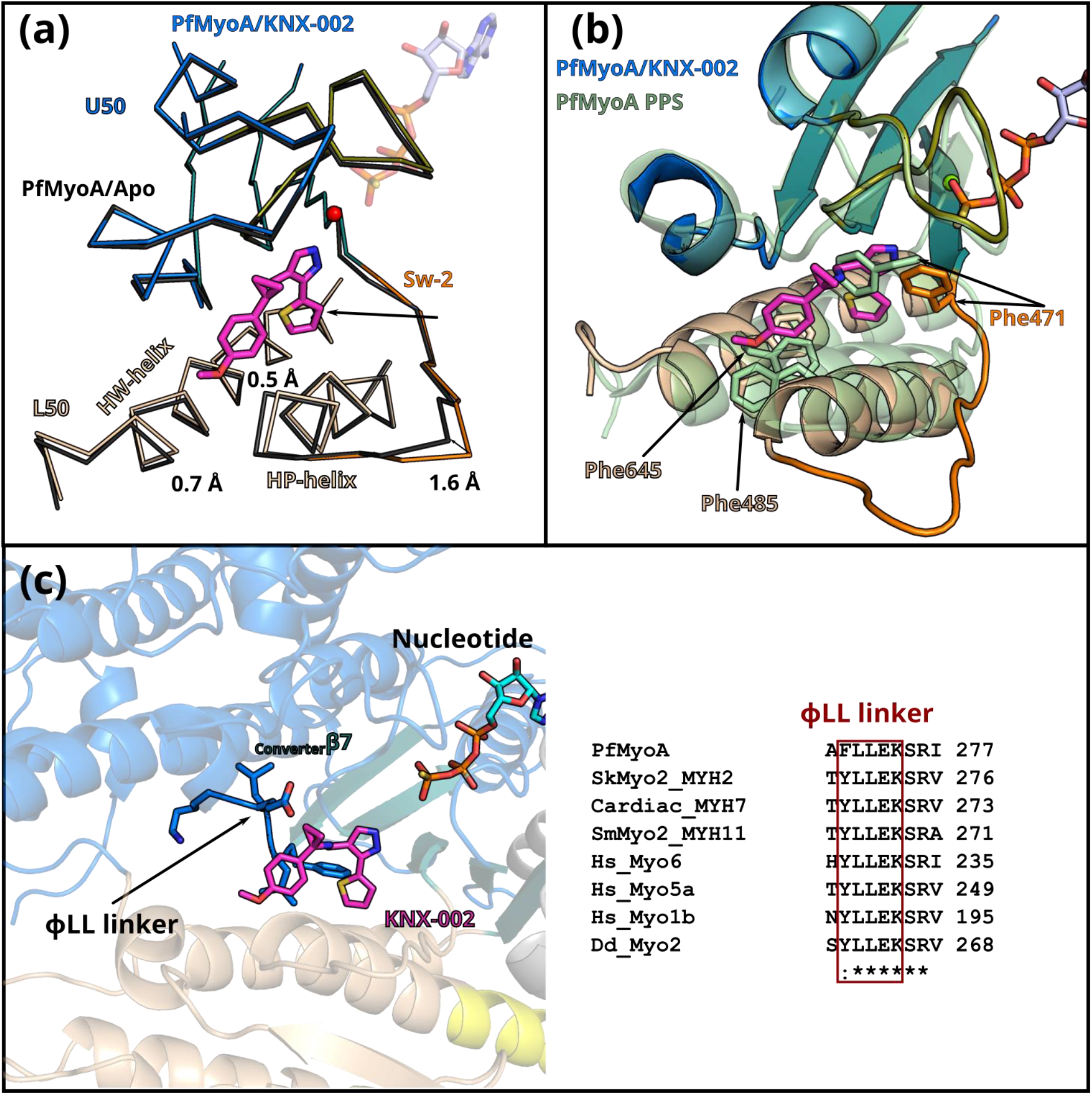
Analysis of the binding pocket of KNX-002. **(a)** Superimposition on the U50 subdomain of PfMyoA bound to KNX-002 (colored by subdomains, PfMyoA/KNX-002) and PfMyoA in the apo condition (black, PfMyoA/Apo). Small local accommodations are induced in the pocket by the binding of the drug, the most remarkable being a shift in Switch-2 (Sw-2) position. **(b)** Superimposition of PfMyoA/KNX-002 and PfMyoA in the pre-powerstroke (PPS) state (PDB code 6YCX,^3^ on the U50 subdomain. The superimposition shows that KNX-002 cannot bind in PPS. In particular, a clash would occur if Switch-2 rearranges to close the back-door as found in PPS. Note in particular, the position F471 adopts in the PPS state. **(c)** Cartoon representation showing the ΦLL linker, a conserved element located after the _Transducer_β7, in the U50. Alignment of myosin sequences show the conservation of the ΦLL linker: PfMyoA; human fast skeletal muscle myosin 2 (SkMyo2_MYH2); human β-cardiac muscle myosin 2 (Cardiac_MYH7); human smooth muscle myosin 2 (SmMyo2_MYH11); human myosin 6 (Hs_Myo6); human myosin 5a (Hs_Myo5a); human myosin-1b (Hs_Myo1b); *Dictyostelium discoidum* myosin 2 (DdMyo2).

**Supplementary Figure 4.**
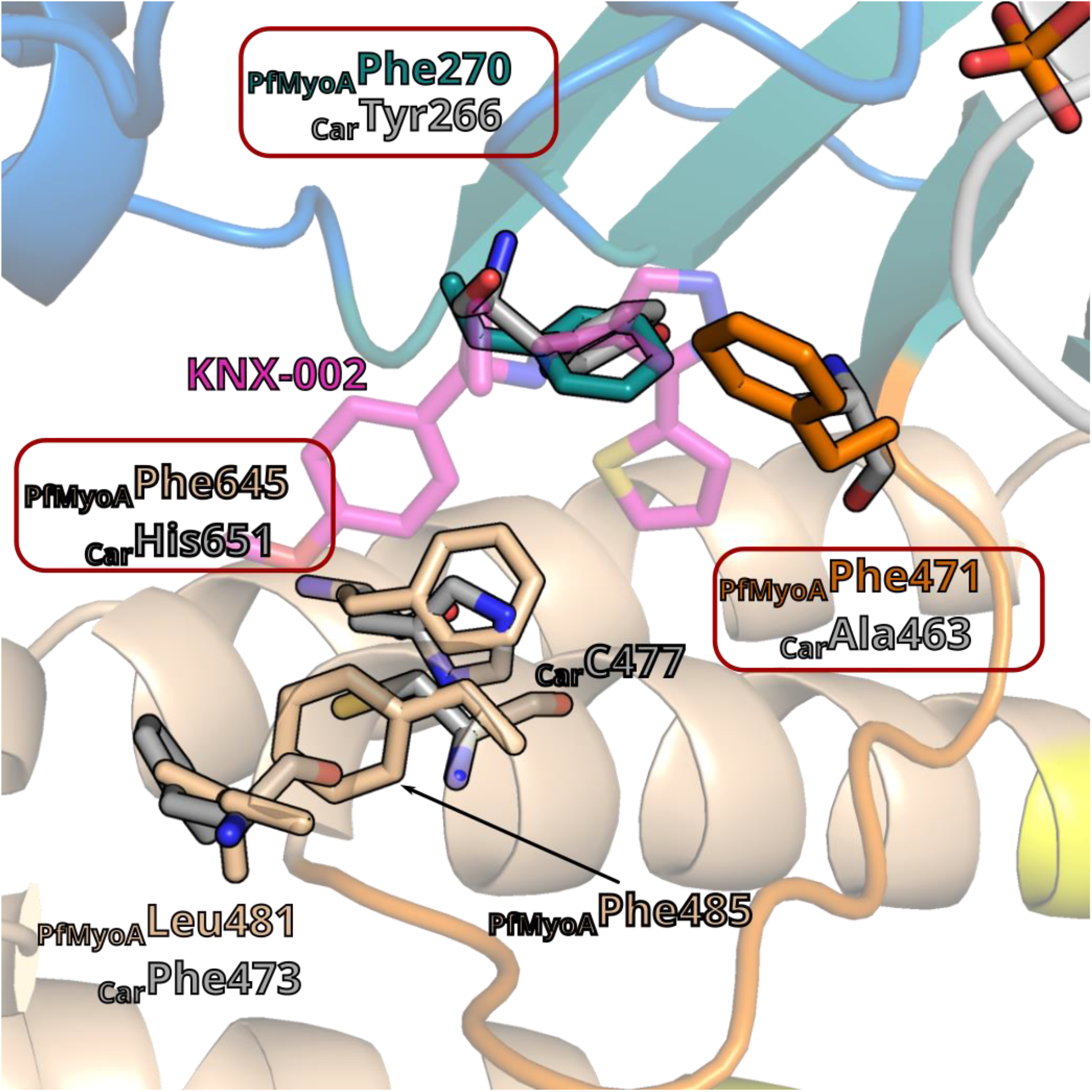
Validation of the KNX-002 binding pocket. Representation of the differences existing in the KNX-002 binding pocket in PfMyoA and in β-cardiac myosin in the PR state (PDB code 6FSA, ^4^, the two structures are aligned on the U50 subdomain of PfMyoA. The sequence differences in cardiac are represented as stick in grey (for detail see **Fig. 3f**). Residues mutated to investigate the pocket are represented in a red box.

**Supplementary Figure 5.**
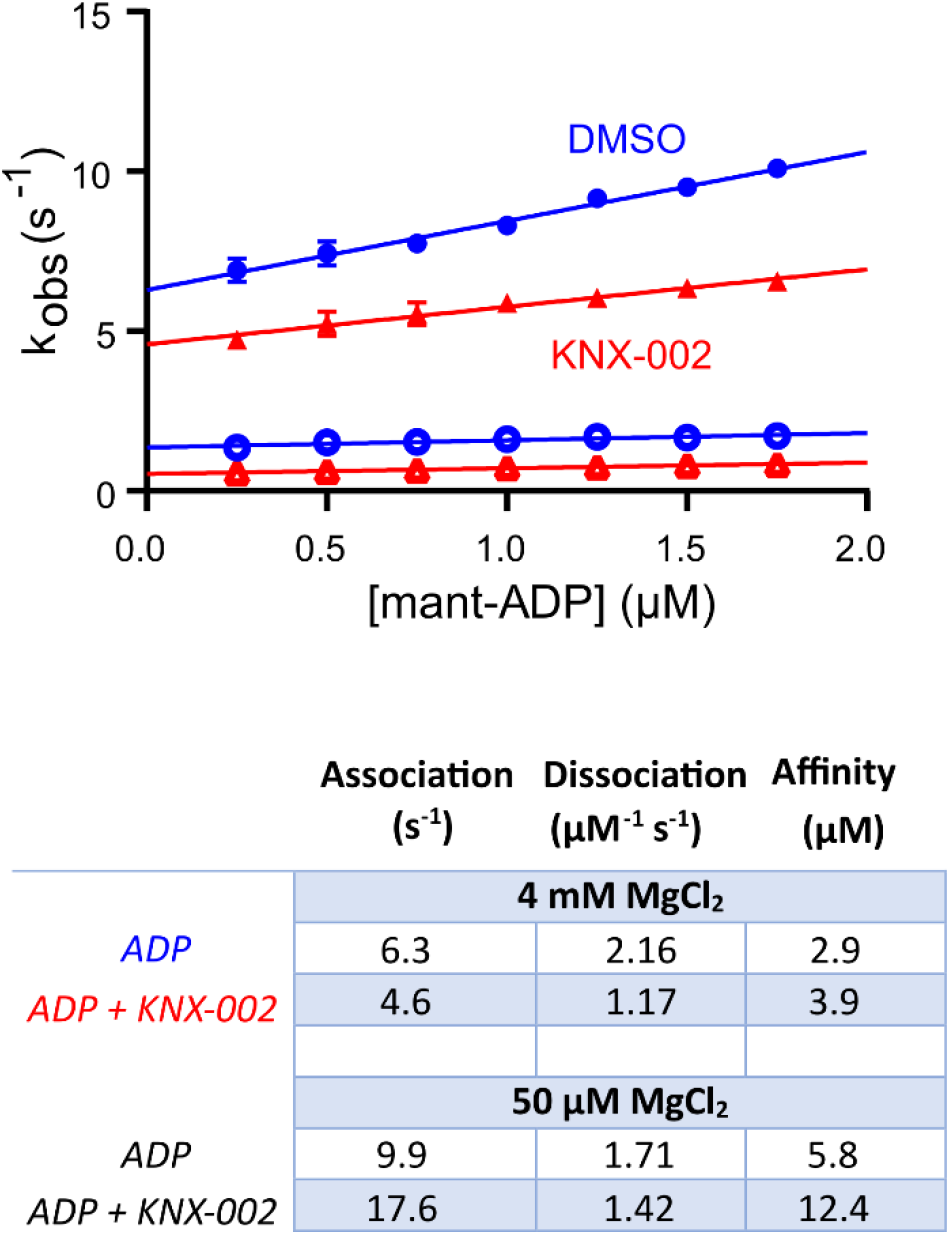
Effect of KNX-002 on the affinity of PfMyoA for ADP. Rates in the absence (blue circles) or presence of 100 μM KNX-002 (red triangles) of the fast (filled symbols) and slow phases (open symbols) from the biphasic transients at 4 mM MgCl_2_ are plotted. The association rate (slope) and dissociation rate (y-intercept) are given in the table below the graph at high and low MgCl_2_ concentration. Data represent two experiments with independent protein preparations.

**Supplementary Figure 6.**
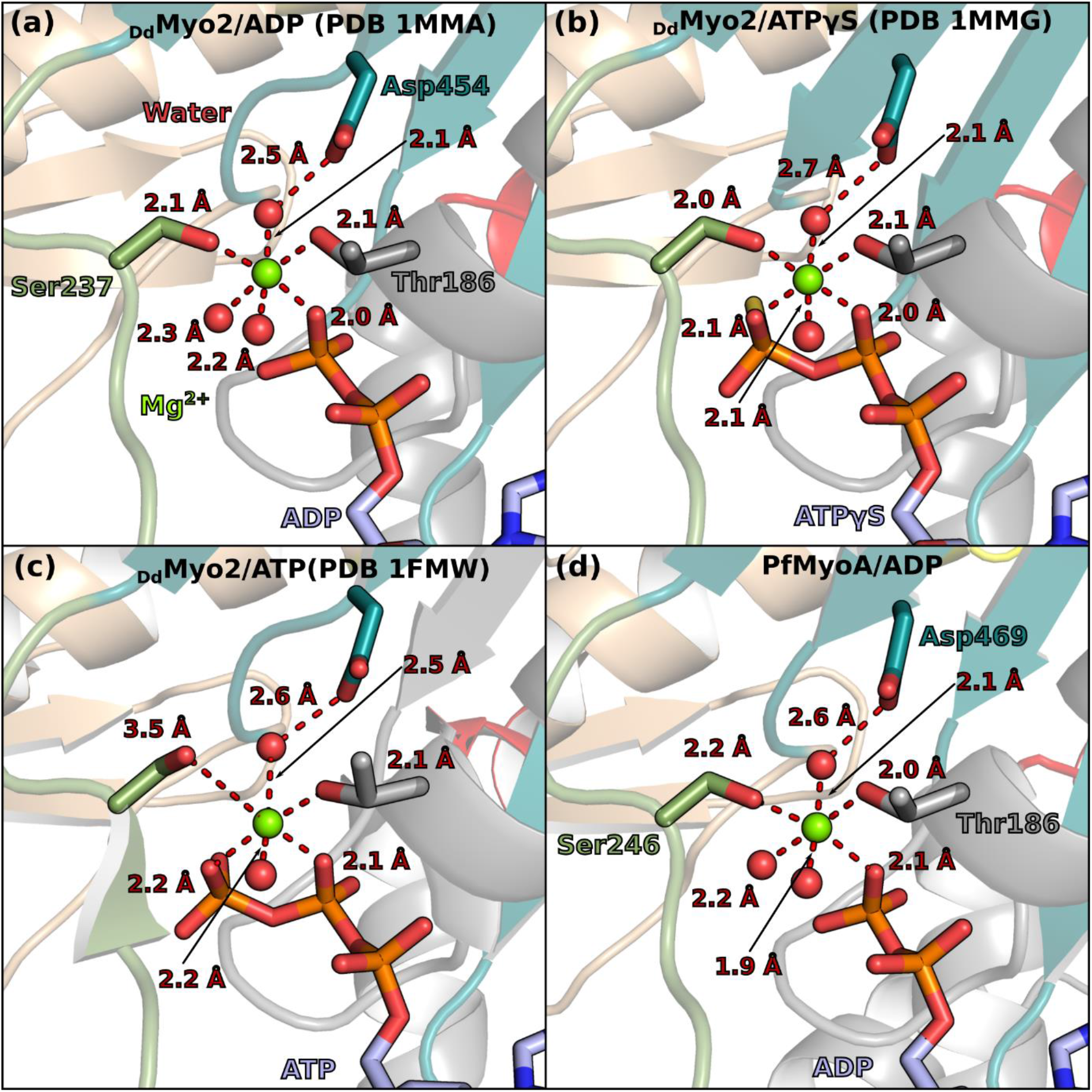
The presence of γ-phosphate does not alter the coordination of Mg^2+^ ion. Comparison of Mg^2+^ coordination in DdMyo2 structure complexed to ADP (PDB code 1MMA), DdMyo2 complexed to ATPγS (PDB code 1MMG) ^5^, **(c)** DdMyo2 structure complexed to ATP (PDB code 1FMW) ^6^. **(d)** For comparison, the coordination of PfMyoA complexed to ADP.

**Supplementary Figure 7.**
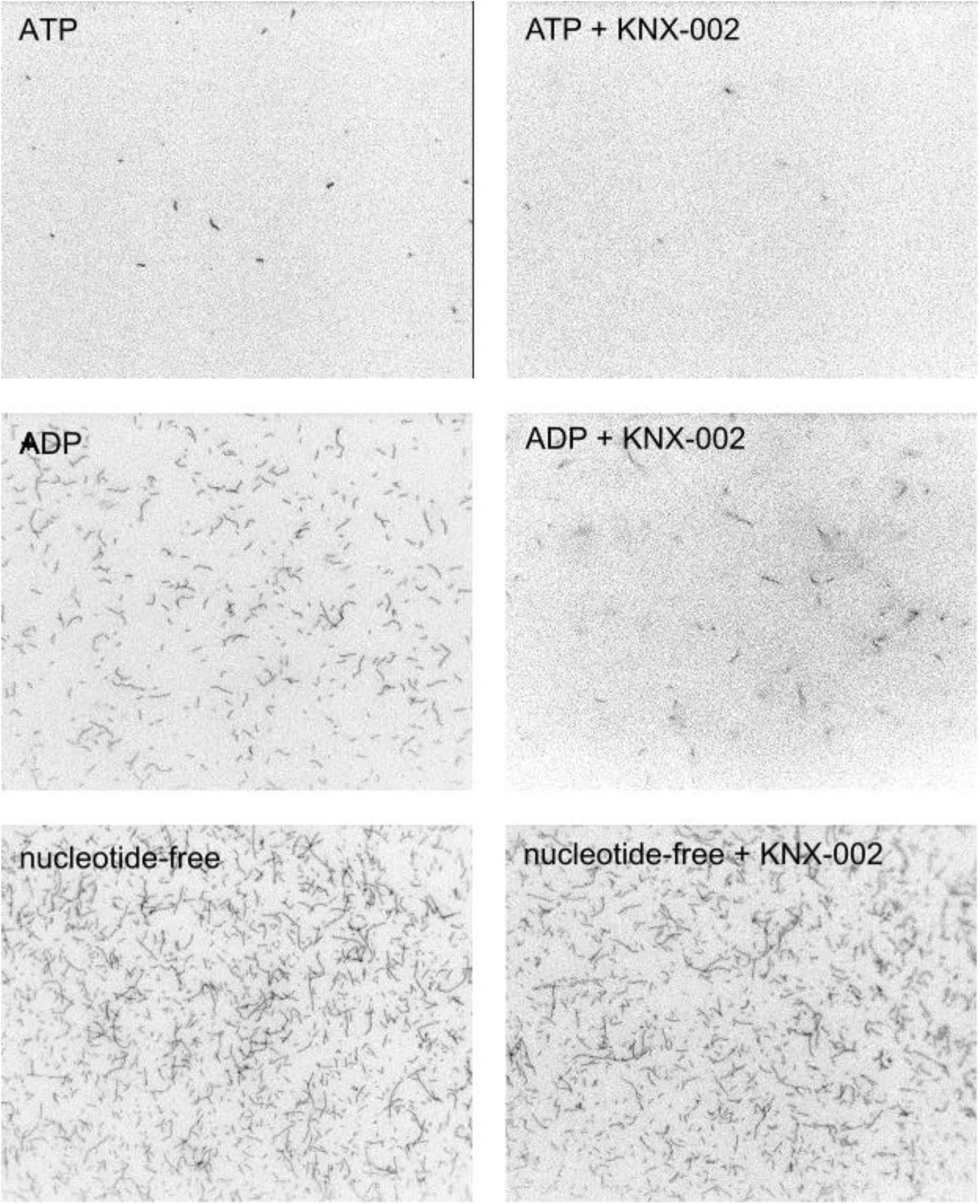
Raw data showing examples of fields of actin bound to surface immobilized PfMyoA. Direct comparison of the amount of actin bound in various nucleotides in the absence or presence of 100 μM KNX-002. Quantitation of the number of bound filaments is shown in **Fig. 5a** and **Supplementary Fig. 8**.

**Supplementary Figure 8.**
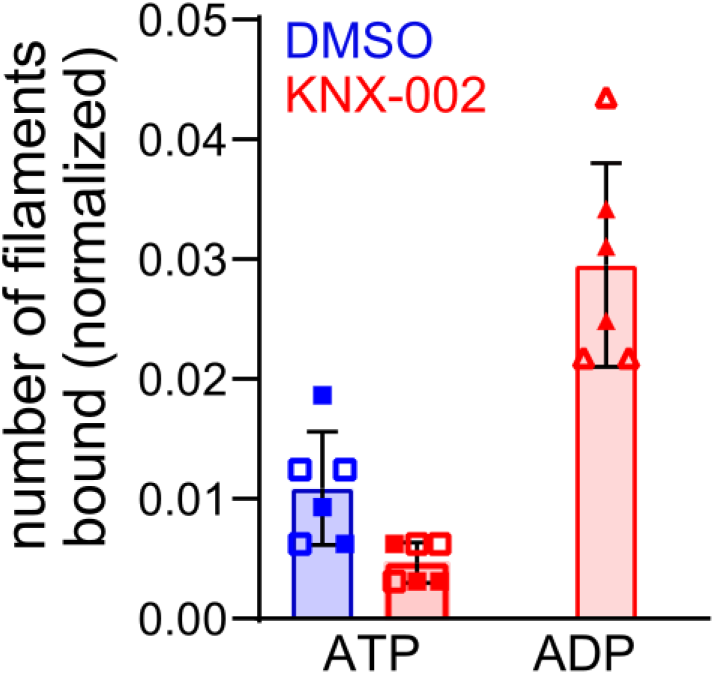
Expanded y-axis for data in **Fig. 5a**. Number of actin filaments bound ± SD to surface immobilized PfMyoA (see Methods): ATP, 0.011 ± 0.004; ATP(+ KNX-002), 0.008 ± 0.030; ADP, 0.59 ± 0.05 (see **Fig. 5a**); ADP(+ KNX-002) 0.030 ± 0.008. Data represent two experiments (open and filled symbols) using independent protein preparations. No significant differences were observed between any pairs of the illustrated data: ATP ± KNX-002 (p>0.999), ADP+KNX-002 and ATP+KNX-002 (p=0.971), ADP+KNX-002 and ATP (p=0.992) (ANOVA followed by Tukey’s post-hoc test). Conditions: 25 mM imidazole pH 7.5, 150 mM KCl, 4 mM MgCl_2_, 1 mM EGTA, 10 mM DTT, 0% methylcellulose, 1% DMSO, 30°C. 2 mM MgATP or MgADP were added for +ATP or +ADP condition, respectively.

**Supplementary Figure 9.**
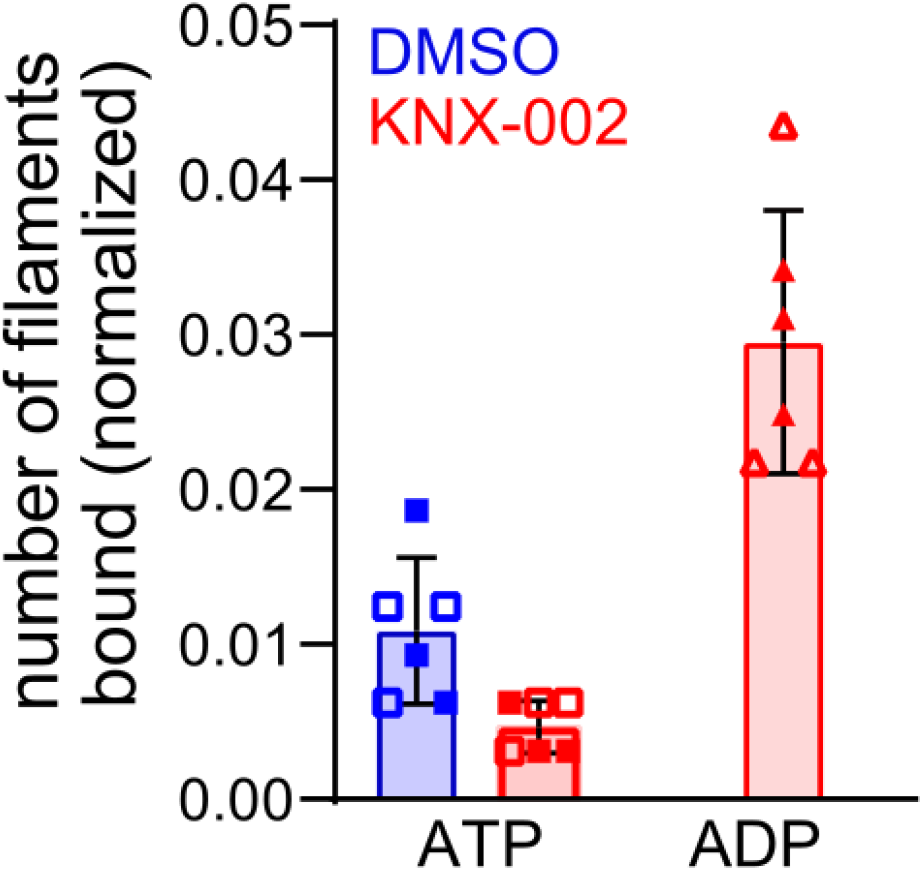
KNX-002 does not affect the rate of ATP induced actomyosin dissociation. 0.3 μM PfMyoA and 0.4 μM skeletal actin were rapidly mixed with ATP in either 1% DMSO (blue open squares) or 100 μM KNX-002 (red filled circles). The apparent second order binding constant ± SD: 1% DMSO, 1.46 ± 0.14 μM^−1^ s^−1^; KNX-002, 1.53 ± 0.33 μM^−1^ s^−1^. Conditions: 10 mM HEPES pH 7.5, 50 mM KCl, 4 mM MgCl_2_, 1 mM EGTA, 1% DMSO, 15°C.

**Supplementary Table 1.** Evaluation of the toxicity of KNX-002 on human cell types: MRC5 (fibroblasts), RPE1 (epithelial cells), HepG2 (hepatic cells). The percentage of cell viability with standard deviation in brackets are represented function of KNX-002 concentration (in μM).

| **Cells type** | **10 $\mu\text{M}$** | **20 $\mu\text{M}$** | **50 $\mu\text{M}$** | **100 $\mu\text{M}$** | **200 $\mu\text{M}$** |
| --- | --- | --- | --- | --- | --- |
| *MRC5* | 99.147 (± 1.702) | 102.566 (± 2.056) | 95.825 (± 2.777) | 82.478 (± 5.889) | 22.642 (± 8.405) |
| *RPE1* | 107.810 (± 1.876) | 109.241 (± 4.010) | 107.961 (± 4.746) | 109.176 (± 1.730) | 0.937 (± 1.321) |
| *HepG2* | 97.119 (± 5.247) | 79.141 (± 2.799) | 46.044 (± 2.830) | 27.380 (± 3.438) | 0.695 (± 0.143) |

**Supplementary Table 2.**
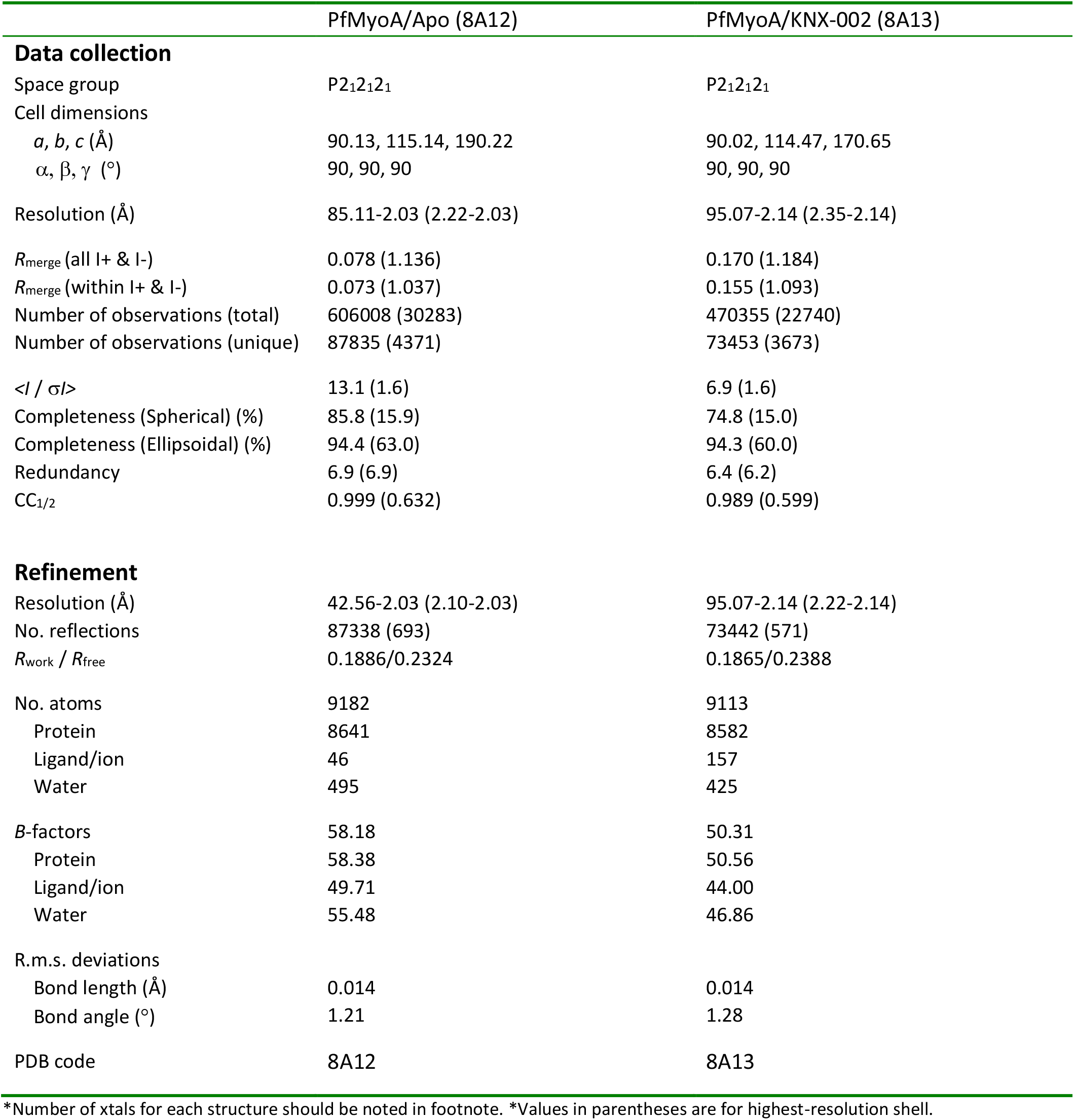
Data collection and refinement statistics (molecular replacement)

|  | PfMyoA/Apo (8A12) | PfMyoA/KNX-002 (8A13) |
| --- | --- | --- |
| **Data collection** |  |  |
| Space group | P2<sub>1</sub>2<sub>1</sub>2<sub>1</sub> | P2<sub>1</sub>2<sub>1</sub>2<sub>1</sub> |
| Cell dimensions |  |  |
| *a*, *b*, *c* (Å) | 90.13, 115.14, 190.22 | 90.02, 114.47, 170.65 |
| $\alpha$, $\beta$, $\gamma$ (°) | 90, 90, 90 | 90, 90, 90 |
| Resolution (Å) | 85.11-2.03 (2.22-2.03) | 95.07-2.14 (2.35-2.14) |
| *R*<sub>merge</sub> (all I+ & I-) | 0.078 (1.136) | 0.170 (1.184) |
| *R*<sub>merge</sub> (within I+ & I-) | 0.073 (1.037) | 0.155 (1.093) |
| Number of observations (total) | 606008 (30283) | 470355 (22740) |
| Number of observations (unique) | 87835 (4371) | 73453 (3673) |
| $\langle I / \sigma I \rangle$ | 13.1 (1.6) | 6.9 (1.6) |
| Completeness (Spherical) (%) | 85.8 (15.9) | 74.8 (15.0) |
| Completeness (Ellipsoidal) (%) | 94.4 (63.0) | 94.3 (60.0) |
| Redundancy | 6.9 (6.9) | 6.4 (6.2) |
| CC<sub>1/2</sub> | 0.999 (0.632) | 0.989 (0.599) |
| **Refinement** |  |  |
| Resolution (Å) | 42.56-2.03 (2.10-2.03) | 95.07-2.14 (2.22-2.14) |
| No. reflections | 87338 (693) | 73442 (571) |
| *R*<sub>work</sub> / *R*<sub>free</sub> | 0.1886/0.2324 | 0.1865/0.2388 |
| No. atoms | 9182 | 9113 |
| Protein | 8641 | 8582 |
| Ligand/ion | 46 | 157 |
| Water | 495 | 425 |
| *B*-factors | 58.18 | 50.31 |
| Protein | 58.38 | 50.56 |
| Ligand/ion | 49.71 | 44.00 |
| Water | 55.48 | 46.86 |
| R.m.s. deviations |  |  |
| Bond length (Å) | 0.014 | 0.014 |
| Bond angle (°) | 1.21 | 1.28 |
| PDB code | 8A12 | 8A13 |

\*Number of xtals for each structure should be noted in footnote. \*Values in parentheses are for highest-resolution shell.

**Supplementary Table 3.** Contacts between ligand and protein residues.

| PfMyoA | KNX-002 (this study) | Dd Myo2 (1YV3) | Blebbistatin |
| --- | --- | --- | --- |
| Hp-hL481 | (D) Methoxyphenyl | Hp-hF466 | Phenyl (D) |
| Hp-hE482 |  | Hp-hE467 |  |
| Hp-hF485 |  | Hp-hC470 |  |
| Hw-hI641 |  | Hw-hV630 |  |
| U50L272 mc |  | U50L263 mc | (D) few interactions |
| Hp-hI486 | (A,D) | Hp-hI471 | (C) |
| U50L271 | (A few, D) | U50L262 | (B,C,D) (electrostatic) |
| U50F270 | (A) Thiophene<br>(pi-stacking) | U50Y261 | (A,B) (pi-stacking) |
| Hp-hT489 | (A) | Hp-hT474 | (A) |
| Hw-hF645 | (A) | Hw-hV634 | (A,B,D) |
| Hw-hQ648 |  | Hw-hQ637 | (A) |
| Hw-hL649 |  | Hw-hL638 |  |
| Hw-hL652 |  | Hw-hL641 |  |
| Sw2I470 | (A,B) | Sw2I455 | (A,B) |
| Sw2F471 | (A,B) (pi-stacking) | Sw2S456 | (B,C) (electrostatic) |
| Sw1S246 Carbonyl | (B) Pyrazole<br>Water-Mg<sup>2+</sup><br>(electrostatic) | Sw1S237 Carbonyl |  |
| TransdF248-G249 mc | (B) | TransdF239-G240 mc | (B,C) (electrostatic) |
| Sw1R247 | (C) cyclopropamine | Sw1R238 | (C) few interactions |
| U50E273 | (C) | U50E264 | (D) |

Blue background – sequences differ between PfMyoA and DdMyo2
Orange background – no interaction

**Supplementary Table 4.** Comparison of the kinetic measurements for KNX-002 and Blebbistatin.

| Blebbistatin | KNX-002 |
| --- | --- |
| **Differences** |  |
| <sup>a</sup> Does not prevent ATP hydrolysis<br>Binds preferentially to ADP.Pi | Diminishes ATP hydrolysis (P<sub>i</sub> burst experiment, Fig. 2d) |
| <sup>a</sup> Does not compete with nucleotide binding | Inhibits ATP binding 10-fold (Fig. 4a) |
| <sup>b</sup> In absence of Mg<sup>2+</sup>, Blebbistatin increases the affinity for ADP | KNX-002 has little effect on ADP affinity at both high and low MgCl<sub>2</sub> concentration (Supplementary Fig. 5) |
| <sup>c</sup> Does not weaken binding of M.ADP to A | Inhibits binding of M.ADP to A (Fig. 5a) |
| <sup>a</sup> Binds weakly to apoM (25 µM) compared to IC<sub>50</sub> for ATPase activity (1.6 µM for basal/0.4 µM for actin activated) | Binds to nucleotide-free M (8.6 µM) and M.ADP (11 µM) with same order of magnitude as the IC<sub>50</sub> ATPase activity (3.6 µM for basal/7.2 µM for actin- activated) (Supplementary Fig. 1 and Fig. 1a) |
| **Similarities** |  |
| <sup>a</sup> Does not interfere with binding of M to A | Does not interfere with binding of M to A (Fig. 5a) |
| <sup>a</sup> Does not alter affinity for ADP at mM MgCl<sub>2</sub> | Does not alter affinity for ADP at mM MgCl<sub>2</sub> (Supplementary Fig. 5) |
| <sup>a</sup> Does not interfere with ATP induced dissociation of AM | Does not interfere with ATP induced dissociation of AM (Supplementary Fig. 9) |
| <sup>a</sup> Blocks myosin heads in a state that binds with low affinity to actin | Blocks myosin heads in a state that binds with low affinity to actin (Fig. 5a) |

<sup>a</sup>Kovacs, M. *et al.* J. Biol Chem 279:25557-63 (2004) <sup>7</sup>
<sup>b</sup>Ewert, W. *et al.* Int. J. Mol Sci. 21:7417 (2020) <sup>8</sup>
<sup>c</sup>Takacs, B. *et al.* PNAS 107:6799-6804 (2010) <sup>9</sup>

*<u>**Supplementary Movie 1**– Movement of actin filaments by PfMyoA in the absence of KNX-002.</u>*

Representative movie showing directed motion of actin filaments powered by PfMyoA. Conditions: 25 mM imidazole pH 7.5, 150 mM KCl, 4 mM MgCl_2_, 1 mM EGTA, 10 mM DTT, 0.15% methylcellulose, 1% DMSO, 30°C.

*<u>**Supplementary Movie 2**– In the presence of KNX-002, no directed movement of actin filaments by PfMyoA is observed.</u>*

In the presence of 200 μM KNX-002, few filaments are seen in the field of view and none exhibit directed motion. Note that the presence of KNX-002 causes photobleaching of the rhodamine-phalloidin labeled actin filaments, resulting in grainier images. Conditions: 25 mM imidazole pH 7.5, 150 mM KCl, 4 mM MgCl_2_, 1 mM EGTA, 10 mM DTT, 0.15% methylcellulose, 1% DMSO, 30°C.

**Supplementary data 1.**
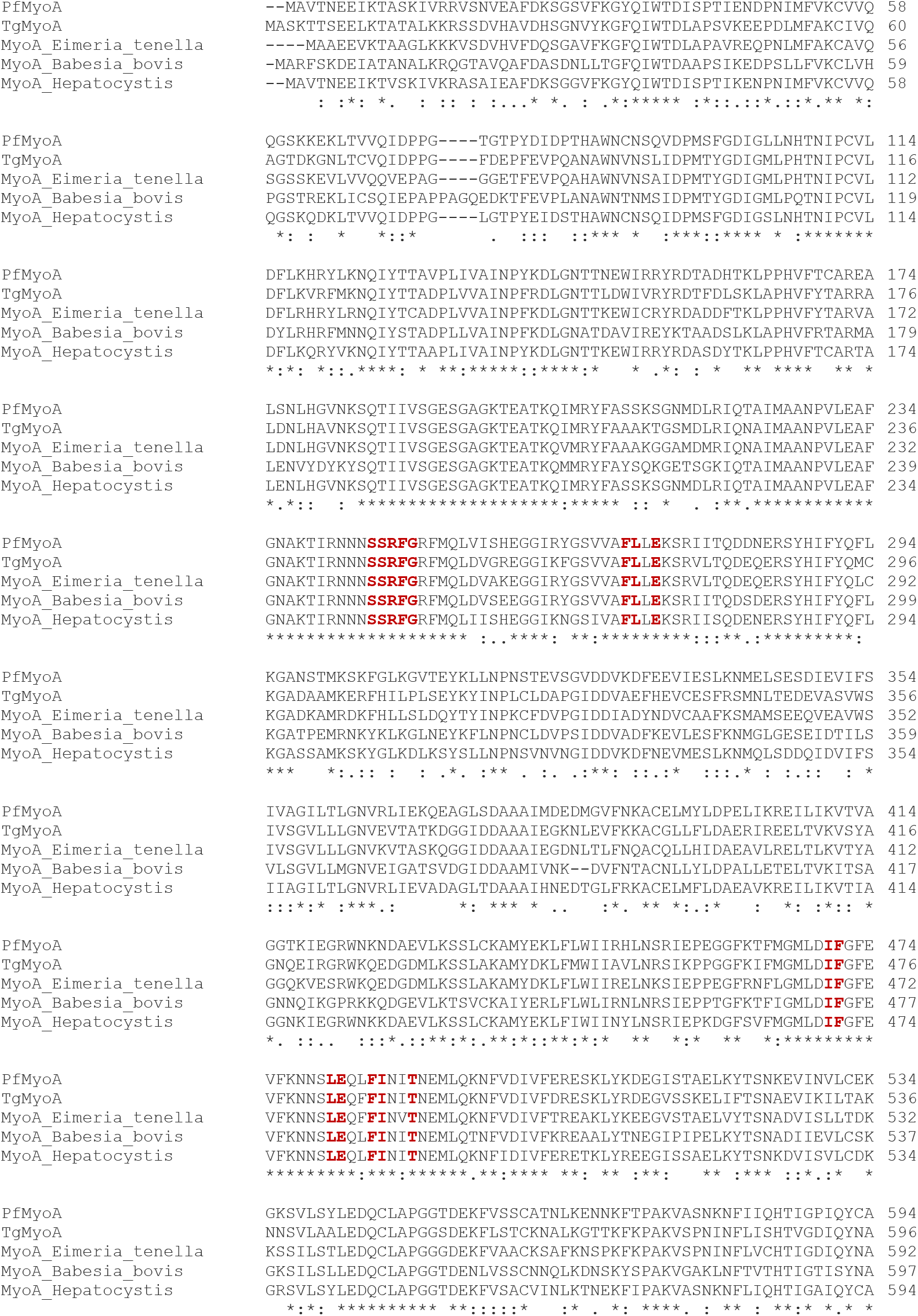
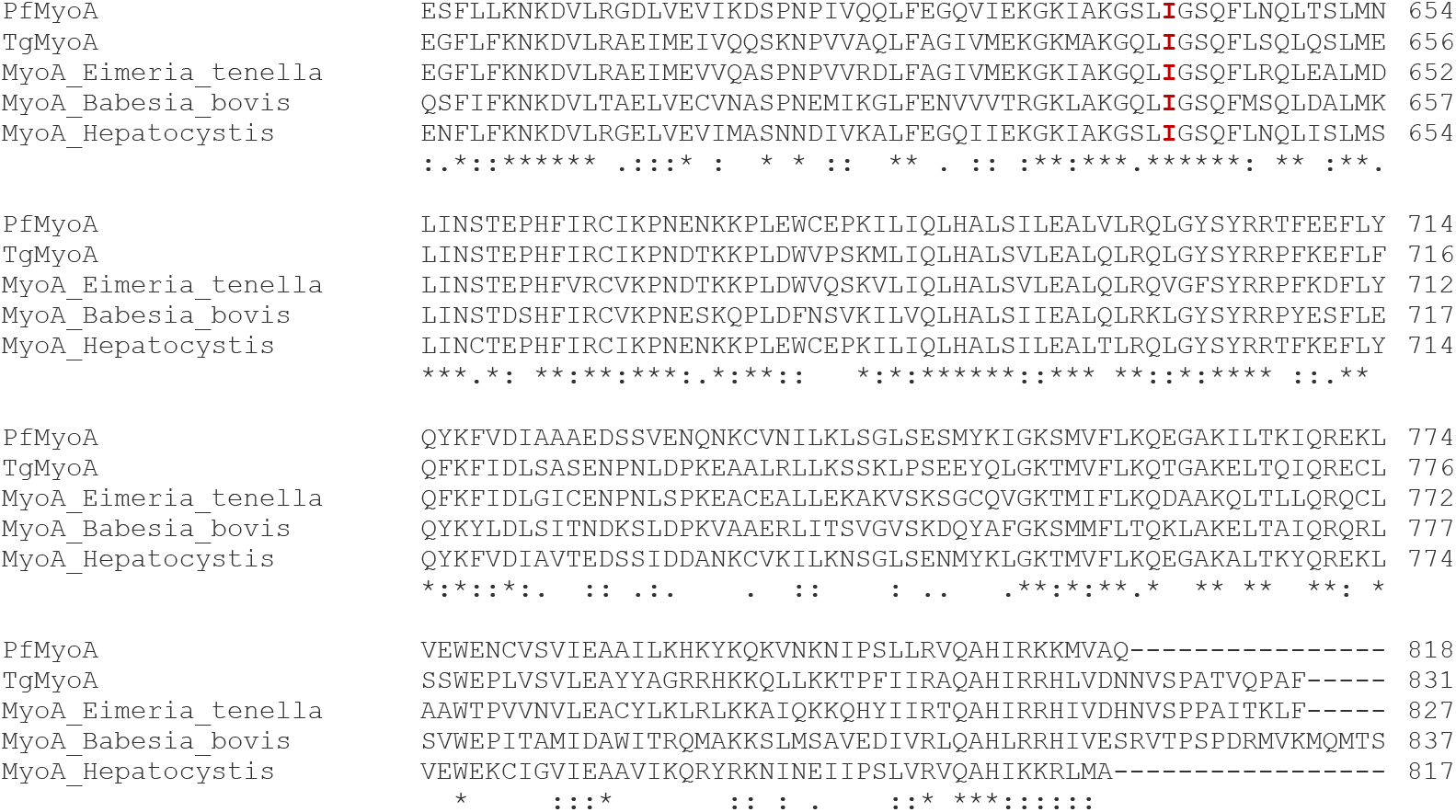
KNX-002 pocket is conserved in sequence in Myosin A from different Apicomplexan genus. Myosin A sequences from different Apicomplexan genus are represented: *Plasmodium falciparum* (PfMyoA); *Toxoplasma gondii* (TgMyoA); *Eimeria tenella*; *Babesia bovis*; *Hepatocystis ssp*.. All the residues involved in KNX-002 binding (in red) are conserved in all the MyoA, indicating that KNX-002 would be able to inhibit all these motors. Fully conserved residues are labelled with a * (asterisk), residues conserved with groups of strongly similar properties are labelled with a: (colon); residues conserved with groups of weakly similar properties are labelled with a. (period). The alignment was realized with ClustalOmega^10^

## References

1. Talman, A. M., Domarle, O., McKenzie, F. E., Ariey, F. & Robert, V. Gametocytogenesis: the puberty of Plasmodium falciparum. Malar. J. 3, 24 (2004).

2. Phillips, M. A. et al. Malaria. Nat. Rev. Dis. Prim. 3, 17050 (2017).

3. Organization, W. H. World malaria report 2021. (World Health Organization).

4. Haldar, K., Bhattacharjee, S. & Safeukui, I. Drug resistance in Plasmodium. Nat. Rev. Microbiol. 16, 156–170 (2018).

5. Balikagala, B. et al. Evidence of Artemisinin-Resistant Malaria in Africa. N. Engl. J. Med. 385, 1163–1171 (2021).

6. Laurens, M. B. RTS,S/AS01 vaccine (Mosquirix™): an overview. Hum. Vaccin. Immunother. 16, 480–489 (2020).

7. Tardieux, I. & Baum, J. Reassessing the mechanics of parasite motility and host-cell invasion. J. Cell Biol. 214, 507–515 (2016).

8. Frenal, K., Dubremetz, J.-F., Lebrun, M. & Soldati-Favre, D. Gliding motility powers invasion and egress in Apicomplexa. Nat. Rev. Microbiol. 15, 645–660 (2017).

9. Robert-Paganin, J. et al. Plasmodium myosin A drives parasite invasion by an atypical force generating mechanism. Nat. Commun. 10, 3286 (2019).

10. Moussaoui, D. et al. Full-length Plasmodium falciparum myosin A and essential light chain PfELC structures provide new anti-malarial targets. Elife 9, (2020).

11. Siden-Kiamos, I. et al. Stage-specific depletion of myosin A supports an essential role in motility of malarial ookinetes. Cell. Microbiol. 13, 1996–2006 (2011).

12. Wall, R. J. et al. Systematic analysis of Plasmodium myosins reveals differential expression, localisation, and function in invasive and proliferative parasite stages. Cell. Microbiol. 21, e13082 (2019).

13. Malik, F. I. et al. Cardiac myosin activation: A potential therapeutic approach for systolic heart failure. Science (80-.). 331, 1439–1443 (2011).

14. Planelles-Herrero, V. J., Hartman, J. J., Robert-Paganin, J., Malik, F. I. & Houdusse, A. Mechanistic and structural basis for activation of cardiac myosin force production by omecamtiv mecarbil. Nat. Commun. 8, 190 (2017).

15. Green, E. M. et al. A small-molecule inhibitor of sarcomere contractility suppresses hypertrophic cardiomyopathy in mice. Science (80-.). 351, 617–621 (2016).

16. Gyimesi, M. et al. Single Residue Variation in Skeletal Muscle Myosin Enables Direct and Selective Drug Targeting for Spasticity and Muscle Stiffness. Cell 183, 335–346.e13 (2020).

17. Sirigu, S. et al. Highly selective inhibition of myosin motors provides the basis of potential therapeutic application. Proc. Natl. Acad. Sci. U. S. A. 113, E7448–E7455 (2016).

18. Rauscher, A. Á., Gyimesi, M., Kovács, M. & Málnási-Csizmadia, A. Targeting Myosin by Blebbistatin Derivatives: Optimization and Pharmacological Potential. Trends Biochem. Sci. 43, 700–713 (2018).

19. Manstein, D. J. & Preller, M. Small Molecule Effectors of Myosin Function. Adv. Exp. Med. Biol. 1239, 61–84 (2020).

20. Day, S. M., Tardiff, J. C. & Ostap, E. M. Myosin modulators: emerging approaches for the treatment of cardiomyopathies and heart failure. J. Clin. Invest. 132, (2022).

21. Utter, M. S., Ryba, D. M., Li, B. H., Wolska, B. M. & Solaro, R. J. Omecamtiv Mecarbil, a Cardiac Myosin Activator, Increases Ca2+ Sensitivity in Myofilaments With a Dilated Cardiomyopathy Mutant Tropomyosin E54K. J. Cardiovasc. Pharmacol. 66, 347–353 (2015).

22. Amgen, Cytokinetics And Servier Announce Start Of METEORIC-HF, The Second Phase 3 Clinical Trial Of Omecamtiv Mecarbil In Patients With Heart Failure. https://wwwext.amgen.com/media/news-releases/2019/02/amgen-cytokinetics-and-servier-announce-start-of-meteorichf-the-second-phase-3-clinical-trial-of-omecamtiv-mecarbil-in-patients-with-heart-failure/.

23. Blake, T. C. A., Haase, S. & Baum, J. Actomyosin forces and the energetics of red blood cell invasion by the malaria parasite Plasmodium falciparum. PLOS Pathog. 16, e1009007 (2020).

24. Yount, R. G., Lawson, D. & Rayment, I. Is myosin a ‘back door’ enzyme? Biophys. J. 68, 44S–47S; discussion 47S-49S (1995).

25. Robert-Paganin, J., Pylypenko, O., Kikuti, C., Sweeney, H. L. & Houdusse, A. Force Generation by Myosin Motors: A Structural Perspective. Chem. Rev. 120, 5–35 (2020).

26. Sweeney, H. L., Houdusse, A. & Robert-Paganin, J. Myosin Structures. Adv. Exp. Med. Biol. 1239, 7–19 (2020).

27. Allingham, J. S., Smith, R. & Rayment, I. The structural basis of blebbistatin inhibition and specificity for myosin II. Nat. Struct. Mol. Biol. 12, 378–379 (2005).

28. Bagshaw, C. R. et al. The magnesium ion-dependent adenosine triphosphatase of myosin. Two-step processes of adenosine triphosphate association and adenosine diphosphate dissociation. Biochem. J. 141, 351–364 (1974).

29. Bagshaw, C. R. & Trentham, D. R. The characterization of myosin-product complexes and of product-release steps during the magnesium ion-dependent adenosine triphosphatase reaction. Biochem. J. 141, 331–349 (1974).

30. Swenson, A. M. et al. Magnesium modulates actin binding and ADP release in myosin motors. J. Biol. Chem. 289, 23977–23991 (2014).

31. Robert-Paganin, J. et al. The actomyosin interface contains an evolutionary conserved core and an ancillary interface involved in specificity. Nat. Commun. 12, 1892 (2021).

32. Kovács, M., Tóth, J., Hetényi, C., Málnási-Csizmadia, A. & Sellers, J. R. Mechanism of blebbistatin inhibition of myosin II. J. Biol. Chem. 279, 35557–35563 (2004).

33. Yahiya, S., Rueda-Zubiaurre, A., Delves, M. J., Fuchter, M. J. & Baum, J. The antimalarial screening landscape-looking beyond the asexual blood stage. Curr. Opin. Chem. Biol. 50, 1–9 (2019).

34. Burrows, J. N. et al. New developments in anti-malarial target candidate and product profiles. Malar. J. 16, 26 (2017).

35. Gyimesi, M. et al. Improved Inhibitory and Absorption, Distribution, Metabolism, Excretion, and Toxicology (ADMET) Properties of Blebbistatin Derivatives Indicate That Blebbistatin Scaffold Is Ideal for drug Development Targeting Myosin-2. J. Pharmacol. Exp. Ther. 376, 358–373 (2021).

36. Bookwalter, C. S. et al. Reconstitution of the core of the malaria parasite glideosome with recombinant Plasmodium class XIV myosin A and Plasmodium actin. J. Biol. Chem. 292, 19290–19303 (2017).

37. Pardee, J. D. & Spudich, J. A. Purification of Muscle Actin. Methods Enzymol. (1982) doi:10.1016/0076-6879(82)85020-9.

38. Henkel, R. D., VandeBerg, J. L. & Walsh, R. A. A microassay for ATPase. Anal. Biochem. 169, 312–318 (1988).

39. McCarthy, A. A. et al. ID30B {--} a versatile beamline for macromolecular crystallography experiments at the ESRF. J. Synchrotron Radiat. 25, 1249–1260 (2018).

40. Kabsch, W. XDS. Acta Crystallogr. Sect. D Biol. Crystallogr. 66, 125–132 (2010).

41. Vonrhein, C. et al. Data processing and analysis with the autoPROC toolbox. Acta Crystallogr. D. Biol. Crystallogr. 67, 293–302 (2011).

42. McCoy, A. J. et al. Phaser crystallographic software. J. Appl. Crystallogr. 40, 658–674 (2007).

43. Moriarty, N. W., Grosse-Kunstleve, R. W. & Adams, P. D. electronic Ligand Builder and Optimization Workbench (eLBOW): a tool for ligand coordinate and restraint generation. Acta Crystallogr. D. Biol. Crystallogr. 65, 1074–1080 (2009).

44. Emsley, P. & Cowtan, K. Coot : model-building tools for molecular graphics. Acta Crystallogr. Sect. D Biol. Crystallogr. 60, 2126–2132 (2004).

45. Bricogne, G. et al. BUSTER version 2.10.2. Cambridge, United Kingdom Glob. Phasing Ltd. (2017).

46. Liebschner, D. et al. Macromolecular structure determination using X-rays, neutrons and electrons: recent developments in Phenix. Acta Crystallogr. Sect. D, Struct. Biol. 75, 861–877 (2019).

47. Berman, H. M. et al. The Protein Data Bank. Nucleic Acids Res. 28, 235–242 (2000).

48. Abraham, M. J. et al. Gromacs: High performance molecular simulations through multi-level parallelism from laptops to supercomputers. SoftwareX 1–2, 19–25 (2015).

49. Jo, S., Kim, T., Iyer, V. G. & Im, W. CHARMM-GUI: A web-based graphical user interface for CHARMM. J. Comput. Chem. 29, 1859–1865 (2008).

50. Essmann, U. et al. A smooth particle mesh Ewald method. J. Chem. Phys. 103, 8577–8593 (1995).

51. Schrödinger, L. L. C. & DeLano, W. PyMOL. (2020).

52. Matthews, H., Deakin, J., Rajab, M., Idris-Usman, M. & Nirmalan, N. J. Investigating antimalarial drug interactions of emetine dihydrochloride hydrate using CalcuSyn-based interactivity calculations. PLoS One 12, e0173303 (2017).

## References

1. Llinas, P. et al. How Actin Initiates the Motor Activity of Myosin. Dev. Cell 33, 401–412 (2015).

2. Robert-Paganin, J., Pylypenko, O., Kikuti, C., Sweeney, H. L. & Houdusse, A. Force Generation by Myosin Motors: A Structural Perspective. Chem. Rev. 120, 5–35 (2020).

3. Moussaoui, D. et al. Full-length Plasmodium falciparum myosin A and essential light chain PfELC structures provide new anti-malarial targets. Elife 9, (2020).

4. Robert-Paganin, J., Auguin, D. & Houdusse, A. Hypertrophic cardiomyopathy disease results from disparate impairments of cardiac myosin function and auto-inhibition. Nat. Commun. 9, 4019 (2018).

5. Gulick, J. et al. Transgenic remodeling of the regulatory myosin light chains in the mammalian heart. Circ. Res. 80, 655–664 (1997).

6. Bauer, C. B., Holden, H. M., Thoden, J. B., Smith, R. & Rayment, I. X-ray structures of the apo and MgATP-bound states of Dictyostelium discoideum myosin motor domain. J. Biol. Chem. 275, 38494–38499 (2000).

7. Kovács, M., Tóth, J., Hetényi, C., Málnási-Csizmadia, A. & Sellers, J. R. Mechanism of blebbistatin inhibition of myosin II. J. Biol. Chem. 279, 35557–35563 (2004).

8. Ewert, W., Franz, P., Tsiavaliaris, G. & Preller, M. Structural and Computational Insights into a Blebbistatin-Bound Myosin•ADP Complex with Characteristics of an ADP-Release Conformation along the Two-Step Myosin Power Stoke. Int. J. Mol. Sci. 21, (2020).

9. Takács, B. et al. Myosin complexed with ADP and blebbistatin reversibly adopts a conformation resembling the start point of the working stroke. Proc. Natl. Acad. Sci. U. S. A. 107, 6799–6804 (2010).

10. McWilliam, H. et al. Analysis Tool Web Services from the EMBL-EBI. Nucleic Acids Res. 41, W597–600 (2013).

